# Cross-competition shapes CD8+ T cell hierarchies and differentiation after RNA vaccination

**DOI:** 10.1101/2025.10.26.684631

**Authors:** Mark J. McCarron, Milena Hornburg, Thomas D. Wu, Ann-Jay Tong, Siri Tahtinen, Martine Darwish, Emily Freund, Ying Feng, David Eisel, Camille-Charlotte Balança, Itai Doron, Huan Lan, Sophie Lehar, Tamaki Nozawa, Yoko Oei, Vincent Javinal, Armando Navarro, Yajun Chestnut, Alan Gutierrez, Sara Wichner, Cecile de la Cruz, Benjamin Haley, Craig Blanchette, Mathias Vormehr, Lena Kranz, Ugur Sahin, Ira Mellman, Jill Schartner, Lélia Delamarre

## Abstract

Immunodominance is a universal feature of adaptive immunity that constrains T cell expansion, clonal diversity and breadth resulting in a narrowly focused T cell response. While observed across diverse priming settings and vaccine platforms, the influence of immunodominance on T cell phenotype remains unclear. Using an mRNA lipoplex vaccine encoding multiple antigens to study how immunodominance influences CD8+ T cell fate, we found that dominant CD8+ T cell responses alter the magnitude and phenotype of subdominant responses through peptide-MHC-I stability-mediated T cell cross-competition. Dominant CD8+ T cell responses preferentially acquired markers associated with terminal differentiation and cytotoxic function, while sub-dominant responses adopted memory-precursor and stem-like features. Removal of dominant responses allowed increased expansion of sub-dominant T cell responses and adoption of terminally differentiated effector phenotypes. These findings reveal that immunodominance dynamically shapes the magnitude, breadth and differentiation of CD8+ T cell responses and highlights opportunities to fine-tune T cell responses for therapeutic vaccination.

## Introduction

Immunodominance describes the tendency of both humoral and cell-mediated immunity to focus the immune response on a limited number of antigens or epitopes^1^. In the case of T cells, favored clones within dominant T cell specificities outcompete and eventually outnumber sub-dominant T cell responses over the lifetime of an immune response. This competitive process helps conserve resources by maintaining a manageable pool of effector cells, but as a result leads to a significant reduction in the diversity of the T cell repertoire^2^, raising important considerations for vaccine strategies aimed at eliciting broad and multispecific T cell responses.

Immunodominance is an often unavoidable and universal feature of T cell biology and has been observed across CD4+ and CD8+ T cell responses^3,4^ and in diverse settings including infectious disease^5^, transplantation^6^, tumor^7^ and vaccination^5^. Factors contributing to this phenomenon include antigen-intrinsic properties such as antigen processing^8^, peptide binding affinity to major histocompatibility complex (MHC) molecules^9^ and stability of peptide-MHC (pMHC) complexes^10,11^, that collectively can influence the abundance and duration of epitope presentation on professional antigen-presenting cells (APCs). Additionally, T cell-intrinsic properties such as homology to self-antigens^12^, the frequency of antigen-specific pre-immune naïve T cells^9^ and T cell receptor (TCR) affinity for pMHC complexes^13^ play key roles. Individually, or collectively, these mechanisms can lead to a competitive suppression of weaker sub-dominant T cell responses by stronger dominant responses that establishes a hierarchy of T cell responses.

Two broad forms of competition between T cells engaging the same APC have been described: (1) inter-T cell competition among T cell clones targeting the same epitope (intra-specificity), and (2) T cell cross-competition among T cells of different specificities (inter-specificity). The first, also termed immunodomination or T cell interference, occurs when dominant T cell clones outcompete other clones targeting the same pMHC-I complex, leading to restricted clonal diversity within the epitope-specific response^5,14^. The second form, T cell cross-competition, refers to competition between T cell clones of distinct specificities, that is recognizing different epitopes presented by the same APC^15,16^. This phenomenon has been observed particularly during periods of intense T cell expansion, such as the boost phase of vaccination^15^ or during secondary immune responses^17^ and is thought to be primarily mediated by limiting antigen presentation that restricts the number of APCs available to stimulate T cell responses^3^. How T cell cross-competition influences the phenotype and effector potential of sub-dominant responses is not understood.

Given the critical role of cytotoxic CD8+ T cells in anti-tumor immunity, efforts to enhance their abundance through vaccination have long been pursued^18^. A decade ago, the discovery that tumor-specific neoantigens, mutated proteins that arise from somatic mutations in cancer cells, are the primary targets of protective anti-tumor immunity sparked renewed efforts and investment in the cancer vaccine field. Therapeutic vaccination strategies designed to elicit T cell responses to tumor neoantigens were initially demonstrated in preclinical models^19–21^, and later validated in human cancer patients^22–27^. Multiple vaccine platforms including peptide-, mRNA-, DNA-, dendritic cell (DC)- and viral vector-based approaches are actively under investigation^28–33^. Among these, RNA-based vaccines have shown particular promise due to their strong immunogenicity and flexibility in encoding individualized, patient-specific neoantigens. Pre-clinical and clinical studies have shown that mRNA-LPX can generate T cell responses of sizable magnitudes^25,34–37^ and durability^38^.

In this study, we investigated the mechanism underlying T cell hierarchy using a clinically relevant mRNA vaccine targeting a range of naturally occurring murine tumor neoantigens, and examined the impact of immunodominance on T cell phenotype. We demonstrate that CD8+ T cell responses organized into reproducible hierarchies governed by T cell cross-competition and in part by pMHC-I stability. We further found that sub-dominant T cells frequently exhibited a less differentiated phenotype, enriched for memory-precursor and stem-like features, while dominant clones were biased toward terminal effector fates. However, in the absence of dominant competitors, sub-dominant T cell responses were capable of adopting effector phenotypes. These data suggest that T cell differentiation is not fixed and determined by the vaccine platform alone, but is instead influenced by the competitive setting. Collectively, these findings reveal that inter-specificity T cell cross-competition can regulate the magnitude and phenotype of T cell responses, with implications for the development of vaccine-induced effector responses.

## Results

### Multi-antigen mRNA lipoplex vaccination remodels CD8+ T cell specificities into a distinct hierarchy

Previously, we identified seven neoantigens presented on MHC-I molecules in the MC38 murine colon adenocarcinoma cell line^39,40^. These included five neoantigens presented by H2-Db (M16/Adpgk, M10/Reps1, M170/Cpne1, M2/Irgq and M111/Aatf) and two by H2-Kb (M152/Dpagt1 and M172/Med12). Since different vaccine platforms have demonstrated differences in immunogenicity for these seven MC38 neoantigens^39,41^, we sought to re-examine their immunogenicity after mRNA vaccination. We used nucleoside-unmodified, uridine-containing mRNA constructs encoding either a single neoantigen (“monotope”) or a polypeptide containing all seven neoantigens (“heptatope”). The mRNA constructs were formulated with liposomes into mRNA lipoplex (RNA-LPX) nanoparticles and administered intravenously to mice^42^. Following three immunizations, an immunization schedule that enables T cell responses to reach a plateau, all seven neoantigens elicited substantial CD8+ T cell responses when delivered individually as monotope RNA-LPX (Fig 1a). In contrast, heptatope RNA-LPX vaccination resulted in a reduced breadth of CD8+ T cell responses which were organized into a different hierarchy (Fig. 1a and Extended data Fig. 1a). Almost all CD8+ T cell responses, except for the M16 response, were substantially reduced in both frequency and total number of T cells (Fig. 1a), with some such as M10 reduced by up to 80-fold (Fig. 1b). This hierarchy was preserved during a memory recall response (Fig. 1c). It was independent of the neoantigen order in the heptatope mRNA, and was maintained when neoantigens were co-delivered on separate mRNAs (Extended Data Fig. 1 b,c). Collectively, these data suggest the development of a stable and reproducible hierarchy of neoantigen-specific CD8+ T cell specificities following multi-neoantigen RNA-LPX vaccination.

**Fig. 1:**
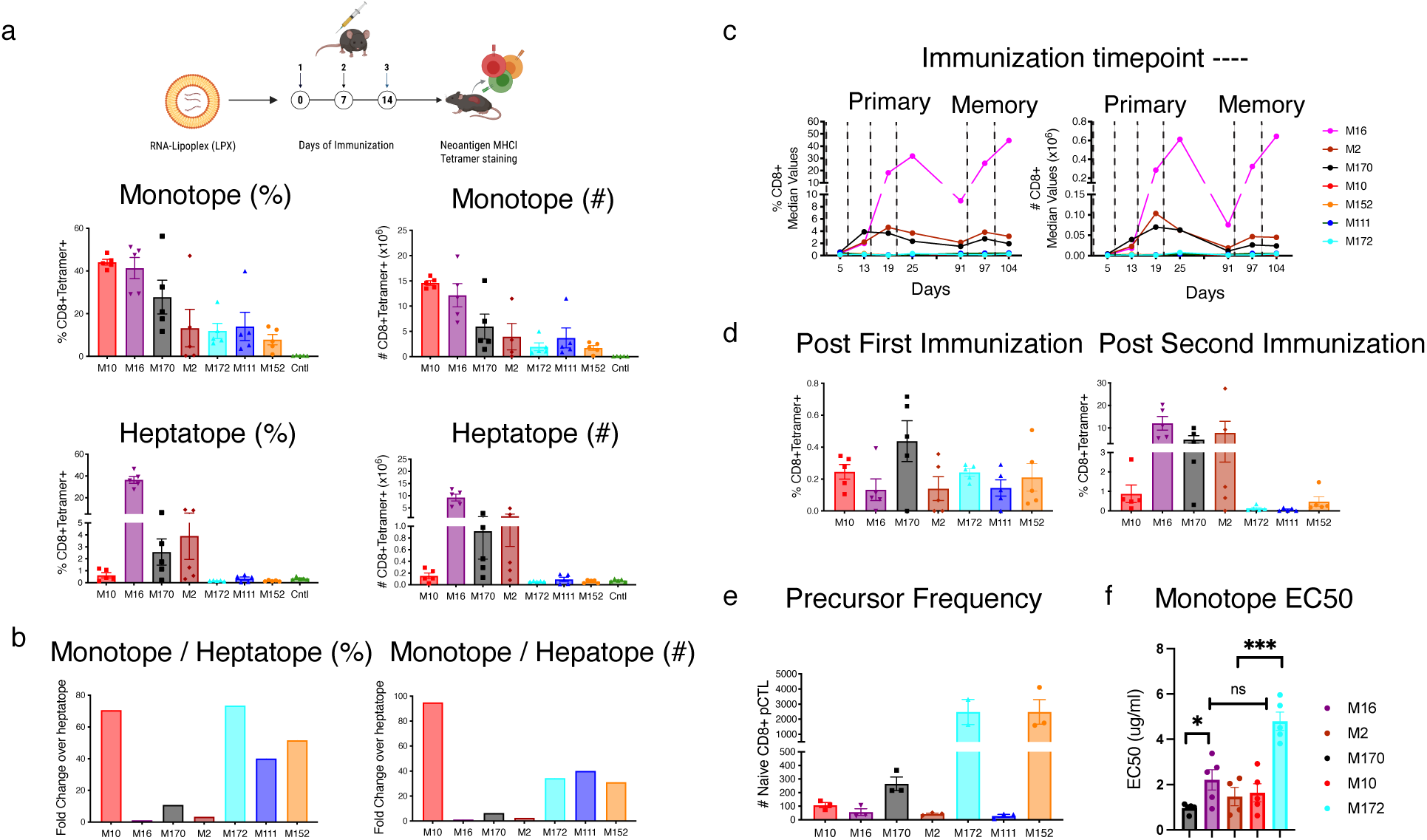
Distinct hierarchy of CD8+ T cell responses following multi-neoantigen RNA-LPX vaccination. a,b C57BL/6 mice were immunized three times on days 0, 7 and 14 with either 50µg of RNA-LPX monotope encoding a single neoantigen or 50µg heptatope RNA-LPX vaccine encoding all 7 neoantigens. On day 19, splenic lymphocytes were quantified by flow cytometry with pMHCI tetramers (specificities indicated on x-axis). a, Neoantigen-specific CD8+ T cell responses as a percentage of total CD8+ T cells and the total number of neoantigen-specific CD8+ T cells (x106) are shown (n=5 individual mice). b, Fold change in the percentage and total number of neoantigen-specific CD8+ T cells following single neoantigen or multi-neoantigen RNA-LPX immunization (monotype/heptatope) is also demonstrated. c, Neoantigen specific CD8+ T cell responses were longitudinally tracked in the peripheral blood following heptatope RNA-LPX immunization. pMHCI tetramer staining during the primary and memory recall responses is shown. Dotted lines indicate the immunization schedule. pMHCI tetramer-stained CD8+ T cells were quantified by flow cytometry and are shown as a percentage of total CD8+ T cells and total number of neoantigen specific T cells. Median of n=10 for primary response, and n=5 for memory response. d, pMHCI tetramer staining was used to quantify neoantigen-specific populations in the blood 5 days after one or two heptatope RNA-LPX immunizations (n=5). e, Naïve precursor CD8+ T cells (pCTL) were enriched from non-immunized mice and quantified. Graph shows the total number of pCTL for each specificity. Each dot represents an individual mouse (n=3). f, C57BL/6 mice were immunized on days 0, 7 and 14 with RNA-LPX monotype vaccines that each encoded a single neoantigen. Functional avidity was assessed by IFNg Elispot assay using titrated peptides. The neoantigen peptide concentration (EC50) at which T cells showed 50% of maximal activity was calculated (n=5). Each dot represents an individual mouse n=5. One-way anova. *p<0.5, **p<0.01, ***p<0.001, ****p<0.0001

### CD8+ T cell hierarchy is established at a time of exponential cell expansion and is independent of the precursor frequency and TCR functional avidity

To determine when the heptatope RNA-LPX hierarchy was established, we analyzed neoantigen-specific CD8+ T cell responses in the blood after each immunization (Fig. 1d). The hierarchy emerged following the second immunization, with the M16 neoantigen becoming the dominant T cell response. These results suggest that the hierarchical ordering of T cell specificities occurs during the phase of exponential CD8+ T cell expansion following boost immunization. Consistent with this, we observed no correlation between the neoantigen-specific T cell precursor frequency and the magnitude of the fully developed response to each neoantigen (Fig. 1e).

Surprisingly, functional avidity measurements of the polyclonal T cell responses after three immunizations with individual monotope RNA-LPX vaccines revealed that the hierarchy is formed independently of TCR functional avidity. While the polyclonal CD8+ T cell response to M172 displayed the lowest avidity (highest EC50), the T cell response to the weaker M10 T cell response exhibited an EC50 comparable to stronger responses (Fig. 1f). Insufficient cell numbers were obtained for reliable EC50 calculations for M152 and M111 following monotope vaccination.

These results suggest that the hierarchy that emerges after the second immunization with the heptatope RNA-LPX vaccine reflects the accumulation of expanded clones and is independent of both TCR functional avidity and precursor frequency.

### CD8+ T cell cross-competition drives the development of the immunodominance hierarchy

The rearrangement of CD8+ T cell responses into a hierarchy occurs following the second immunization, a period marked by substantial T cell expansion. We investigated whether immunodominance driven by T cell cross-competition between different CD8+ T cell specificities contributed to this hierarchy.

T cell cross-competition is defined experimentally as an enhanced response to subdominant determinants when responses to immunodominant determinants are diminished^5,14^. To explore this, we generated a construct in which the most immunodominant neoantigen, M16, was replaced with its non-immunogenic wild-type (WT) counterpart (“heptatope1WT”). Although both the M16 and the WT epitope are processed and presented on MHC-I, the WT epitope is not immunogenic following peptide^39^ or mRNA immunization (data not shown). Removal of the M16 neoantigen resulted in a significant 2-to-3-fold increase in CD8+ T cell responses against most remaining neoantigens, including the other strong M2 response (Fig. 2a). In a second construct (“heptatope3WT”), replacing the three strongest T cell responses (M16, M2, and M170) with their non-immunogenic WT counterparts led to a larger expansion of previously subdominant T cell responses, including up to a 9-fold increase in M10-specific CD8+ T cells (Fig. 2b). These findings demonstrate that the formation of immunodominance hierarchies among neoantigen-specific CD8+ T cell responses is driven by T cell cross-competition where dominant competitors suppress other T cell specificities.

**Fig. 2:**
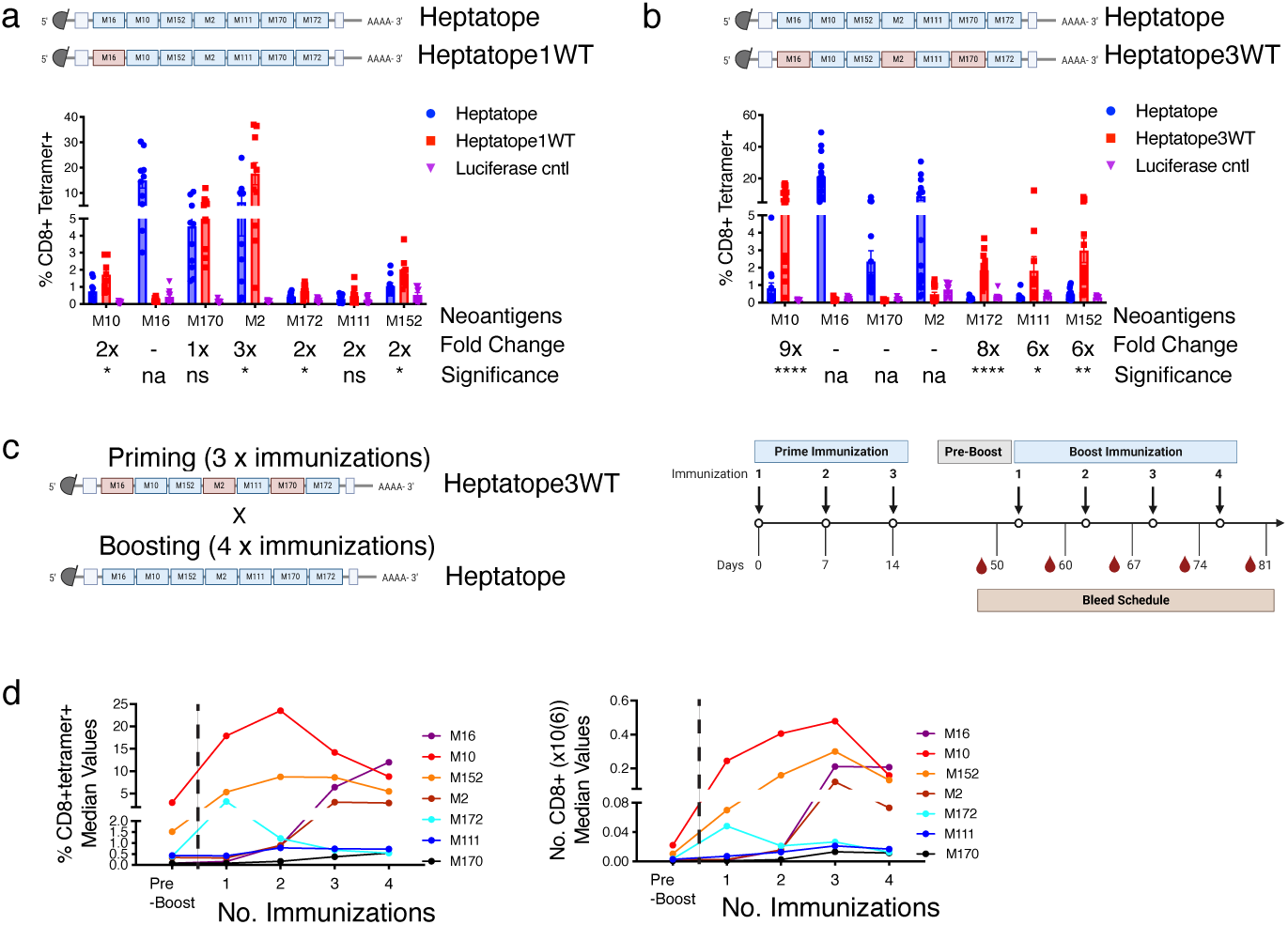
Immunodominance is mediated by T cell cross-competition. a-b, C57BL/6 mice were immunized three times on days 0, 7 and 14 with either RNA-LPX heptatope vaccine encoding encoding all 7 neoantigens or vaccine constructs in which a single neoantigen (Heptatope1WT) or three neoantigens (Heptatope3WT) were replaced by their non-immunogenic, non-neoantigen wild-type counterparts. Flow cytometric quantification of splenic CD8+ T cells was performed with pMHC tetramers (specificities indicated on x-axis) on day 19 (n=10-15). Fold change and statistical significance is denoted below the neoantigen name. Unpaired, two-tailed t-test. ns; not significant (P>0.5), *p<0.5, **p<0.01, ***p<0.001, ****p<0.0001. na; not applicable. c, Experimental design. d, C57BL/6 mice were immunized three times on days 0, 7 and 14 with heptatope3WT RNA-LPX. After a rest period of 40 days, mice were boost immunized with the RNA-LPX heptatope weekly over four weeks. pMHC-I tetramers were used to quantify neoantigen-specific populations in the blood 5 days after each immunization.

We confirmed that T cell hierarchy is dynamically shaped by the properties of concurrent T cell responses by evaluating the immunogenicity of a neoantigen found in the B16F10 melanoma model, Arvcf, in different contexts (Extended Data Fig. 1d). Indeed, Arvcf was dominant when co-immunized with the Nsun2 B16F10 neoantigen, but became subdominant and exhibited reduced immunogenicity when delivered alongside the strong model antigen SIINFEKL (Extended Data Fig. 1d, e).

To further dissect the mechanism underlying cross-competition, we assessed the role of cytokine signaling in this process by combining heptatope RNA-LPX immunization with daily systemic IL-2, a cytokine essential for T cell proliferation and survival. This combination neither increased the total number of neoantigen-specific CD8+ T cells nor altered the immunodominance hierarchy, suggesting that competition for IL-2 is not driving immunodominance (Extended data Fig. 2a). This result was confirmed with a half-life extended IL-2 mRNA which increased CD8+ T cell expansion^43^, without altering T cell hierarchy (Extended data Fig 2b).

Furthermore, we examined the contribution of CD4+ T cell help by incorporating the highly immunogenic CD4 epitope, 2W1S^44^, into the heptatope mRNA (“2W1S octatope mRNA”). 2W1S generated a generous CD4+ T cell response after immunization with 2W1S octatope RNA-LPX (Extended data Fig. 2c), yet did not impact the magnitude or diversity of vaccine-induced CD8+ T cell responses (Extended data Fig. 2d).

One of the earliest observations of immunodominance was the finding that memory CD8+ T cells can suppress the priming of naive CD8+ T cells, driven by the numerical advantage of the recall response which allows these cells earlier access to APCs^45^. We next examined whether RNA-LPX-induced dominant T cell responses can displace established T cell responses. To address this question, mice were first immunized with the heptatope3WT RNA-LPX, which induces T cell responses against M10, M111, M152, and M172. After a forty day rest period, the mice were immunized with heptatope RNA-LPX encoding the additional M16, M2 and M170 neoantigens, and T cell responses were monitored in the blood throughout the immunization protocol (Fig. 2c). Interestingly, following the third immunization with heptatope RNA-LPX, we observed that de novo generation of M16 and M2-specific CD8+ T cell responses was associated with a contraction of the pre-existing M10 and M152-specific CD8+ T cell responses (Fig. 2d). These results suggest that dominant CD8+ T cell responses may suppress pre-existing T cell responses. Furthermore, these data suggest that immunodominance is not simply driven by a T cell numerical advantage, whether following vaccine-mediated expansion (Fig. 2d) or in pre-immune precursor frequency (Fig. 1e).

Immunodominance has also been observed among CD4+ T cell responses^10,46–48^, including that mediated by CD4+ T cell cross-competition^49^. To investigate whether this occurs following RNA-LPX immunization, we generated mRNAs encoding a polypeptide containing six neoantigens from the MC38 tumor model that we found generate exclusively CD4+ T cell responses^50^, plus the 2W1S CD4+ T cell epitope or a non-immunogenic antigen as control. CD4+ T cell responses were unaltered in the presence of a dominant 2W1S response (Extended data Fig. 2e), suggesting that T cell cross-competition was not limiting the magnitude or breadth of CD4+ T cell responses, possibly due to the reduced expansion of CD4+ T cells in comparison to CD8+ T cells. Interestingly, when we co-immunized mice with the strong CD8+ T cell model antigen, SIINFEKL derived from ovalbumin, and M30, a CD4+ T cell neoantigen^37^, we observed a reduction in the CD4+ T cell response (Extended data Fig. 2f), suggesting that CD8+ T cells may cross-compete with CD4+ T cell responses.

### Immunodominance is partially governed by pMHC-I stability rather than competition between neoantigens for MHC-I loading

Next, we investigated whether antigen presentation to T cells influences immunogenicity and immunodominance following RNA-LPX immunization. RNA-LPX is preferentially delivered to splenic DCs where it is translated into a polypeptide^36^. This polypeptide is rapidly processed by the proteasome and the resulting peptides are translocated to the endoplasmic reticulum for loading onto MHC-I. The level and duration of antigen presentation by MHC-I are critical factors in determining the development of T cell responses. Antigen competition for loading onto MHC-I, affinity of the antigen peptide for MHC-I, as well as stability of the pMHC-I complex can all influence antigen presentation by DCs.

We first investigated whether competition between neoantigens for MHC-I binding affects immunogenicity. Of the seven neoantigens evaluated, five bind H2-Db and two bind H2-Kb^39^. We reasoned that competition might only occur for peptides that bind the same allele. To test this, we generated heptatope RNA-LPX vaccines encoding a polypeptide containing one copy of an immunogenic antigen and six copies of a non-immunogenic wild-type antigen, either binding the same allele or a different allele. We selected M16WT to assess competition for H2-Db loading, and M152WT to assess competition for H2-Kb loading, both of which we previously found to be naturally processed and presented on MHC-I with high predicted binding affinity^39^. We experimentally confirmed that binding affinity was relatively similar between WT and mutant neoantigen peptides (Extended Data Fig. 3a). Immunization with a non-competitive M10:M152WT or competitive M10:M16WT RNA-LPX vaccine resulted in equivalent H2-Db M10-specific CD8+ T cell responses (Fig. 3a, left panel). Similarly, the CD8+ T cell response against the H2-Kb M172 neoepitope was unchanged following immunization with a non-competitive M172:M16WT or competitive M172:M152WT RNA-LPX (Fig. 3a, right panel). These findings suggest that competition between neoantigens for access to MHC-I loading is not a limiting factor for immunogenicity and does not contribute to the immunodominance hierarchy in the context of RNA-LPX vaccination.

**Fig. 3:**
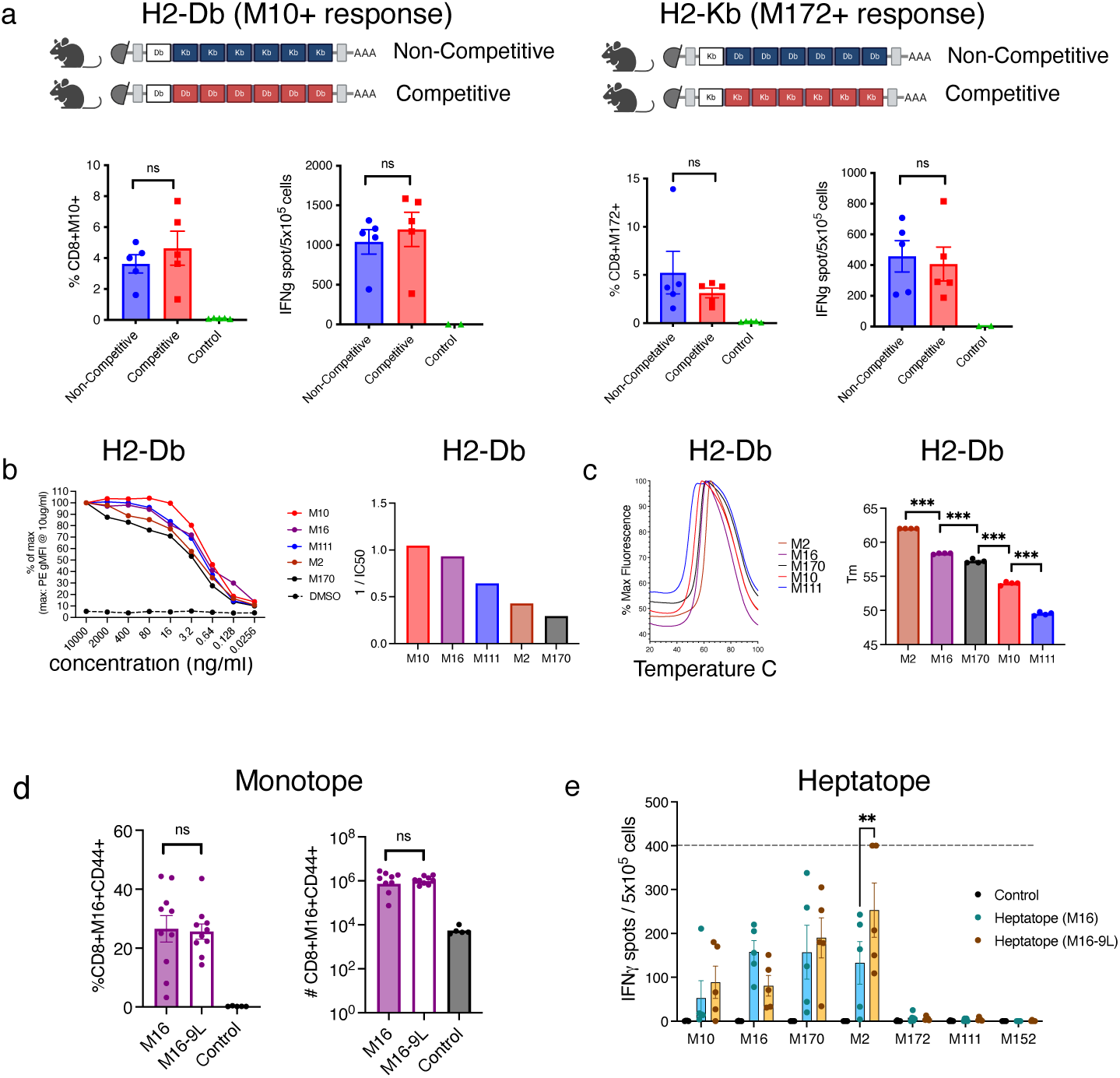
Neoantigen-MHC class I stability partially determines immunodominance. a, C57BL/6 mice were immunized three times on days 0, 7 and 14. To assess competition for H2-Db loading, mice were immunized with either a non-competitive heptatope RNA-LPX vaccine construct encoding M10 and 6 copies of M152WT or a competitive construct encoding M10 and 6 copies of M16WT. To assess competition for H2-Kb loading, mice were immunized with either a non-competitive heptatope RNA-LPX vaccine construct encoding M172 with 6 copies of M16WT or a competitive construct encoding M172 with 6 copies of M152WT. Neoantigen-specific CD8+ T cell responses were measured using two independent assays, pMHCI tetramer+ cells as a percentage of total CD8+ T cells or IFNg spot count per 105 splenocytes. Immunization with RNA-LPX encoding F-luciferase was used as a control. n=5. b, pMHC affinity for each neoantigen was assessed by flow cytometric quantification of H2-Db stabilization on TAP-deficient EL4 cells. Percent of maximum response over a range of neoantigen concentrations is shown and the graph represents the calculated 1/EC50 for each specificity. Data is representative of three individual experiments. c, pMHC complex stability (Koff) for each H2-Db neoantigen was measured by differential scanning fluorimetry following thermal denaturation of soluble pMHC-I complexes. Melting curves were normalized to the minimum and maximum fluorescence values and Tm was calculated using the negative first derivative of RFU values over temperature. n=3 independent studies. d-e, C57BL/6 mice were immunized three times on days 0, 7 and 14 with either 50µg of RNA-LPX monotope encoding a single neoantigen (M16 or M16-9L) or 50µg heptatope RNA-LPX vaccine encoding either the M16 neoantigen (heptatope (M16)) or a mutated M16 variant (Heptatope (M16-9L)) with the other 6 neoantigens. d, On day 19, splenic lymphocytes were quantified by flow cytometry with pMHCI tetramers. Neoantigen-specific CD8+ T cell responses as a percentage of total CD8+ T cells and the total number of neoantigen-specific CD8+ T cells are shown (n=5-10 individual mice). e, Neoantigen-specific CD8+ T cell responses following were measured using IFNg spot count per 105 splenocytes. The dotted line indicates the maximum number of spots that can be counted. Immunization with RNA-LPX encoding F-luciferase was used as a control. Unpaired, two-tailed t-test or two-way anova. ns; not significant (P>0.5), **p<0.01, ***p<0.001.

Next, we generated Tap1-deficient EL4 cell lines overexpressing either the H2-Db or H2-Kb alleles to evaluate the binding affinity of the neoantigens to MHC-I, by quantifying pMHC-I complex that stabilized on the cell surface by flow cytometry. This approach enables comparison of antigens binding the same allele (Fig. 3b and Extended data Fig. 3b). We found that T cell expansion overall reflected antigen/MHC-I binding affinity following monotope RNA-LPX immunization for the H2-Db neoantigens whereby the M10 and M16 neoantigens respectively showed both highest T cell expansion and neoantigen binding affinity (Fig. 3b & Fig. 1a). M170 demonstrated a low affinity in spite of high T cell expansion, which may reflect compensation by a high naïve T cell precursor frequency (Fig. 1e). However, no correlation was observed between the antigen/MHC-I binding affinity and the immunodominance hierarchy following heptatope RNA-LPX immunization (Fig. 3b & Fig. 1a).

It has been shown that pMHC-I stability can be a better predictor of immunogenicity than binding affinity^51–53^. To assess pMHC-I stability, we used differential scanning fluorimetry (DSF) to measure pMHC-I kinetic stability through the thermal denaturation of soluble pMHC-I complexes^54^. As with the affinity assays, this analysis can only compare peptides binding the same allele. For the H-2Db allele, we found that the stronger neoantigens M2, M16 and M170 exhibited higher thermal stability than the less dominant neoantigens (Fig. 3c). A similar trend was observed for H-2Kb, although there were only 2 neoantigens available for analysis (Extended data Fig. 3c). To directly test whether pMHC-I stability can influence T cell cross competition, we produced a less stable M16/MHC-I complex, by introducing a mutation at position 9 of M16, termed M16-9L, that resulted in a substantial ≈5°C reduction in stability (Extended data Fig. 3d). Although M16-9L elicited comparable immunogenicity to parental M16 when delivered as a monotope vaccine (Fig 3d), M16-9L lost its dominant position in the heptatope setting (Fig 3e), enabling expansion of the other responses, most notably a significant increase in the M2-specific CD8+ T cell response. These findings demonstrate that pMHC-I stability is critical to shaping the immunodominance hierarchy.

We validated the totality of these findings using an independent RNA-LPX encoding a polypeptide containing ten neoantigens which generated strong (M16), intermediate (M143) and weak (M86) CD8+ T cell responses (Extended data Fig. 3e-j). Similar to the heptatope RNA-LPX vaccine, reorganization of the neoantigen-specific CD8+ T cell hierarchy was driven by immunodominance (Extended data Fig. 3e,f), independently of the naïve T cell precursor frequency (Extended data Fig. 3g). Moreover, the immunodominance hierarchy mirrored the stability of the pMHC-I complexes with M16 being the strongest and most stably presented neoantigen (Extended data Fig. 3h). For these neoantigens, peptide/MHC-I binding affinity was also correlated with hierarchy (Extended data Fig. 3i,j).

Overall, our data suggest that pMHC-I stability broadly correlates with the observed immunodominance hierarchy, consistent with a recent study showing that endogenous tumor-reactive T cell responses are hierarchical and driven by pMHC-I stability^53^.

### Increasing antigen expression can mitigate immunodominance hierarchies

We reasoned that low pMHC-I stability could be compensated for by increasing antigen expression and presentation. Increasing RNA-LPX dose increases both antigen expression by DCs and their maturation (Fig. 4a), resulting in a corresponding enhancement in both the magnitude and breadth of CD8+ T cell responses (Fig. 4b). Only the dominant M16 and M2 T cell responses were detectable at lower vaccine doses (2-10µg), though non-significantly increased (Extended data Fig.4a). The remaining weaker responses were detectable only at the highest 50µg vaccine dose tested, suggesting that the vaccine dose threshold for priming varies between neoantigens.

**Fig. 4:**
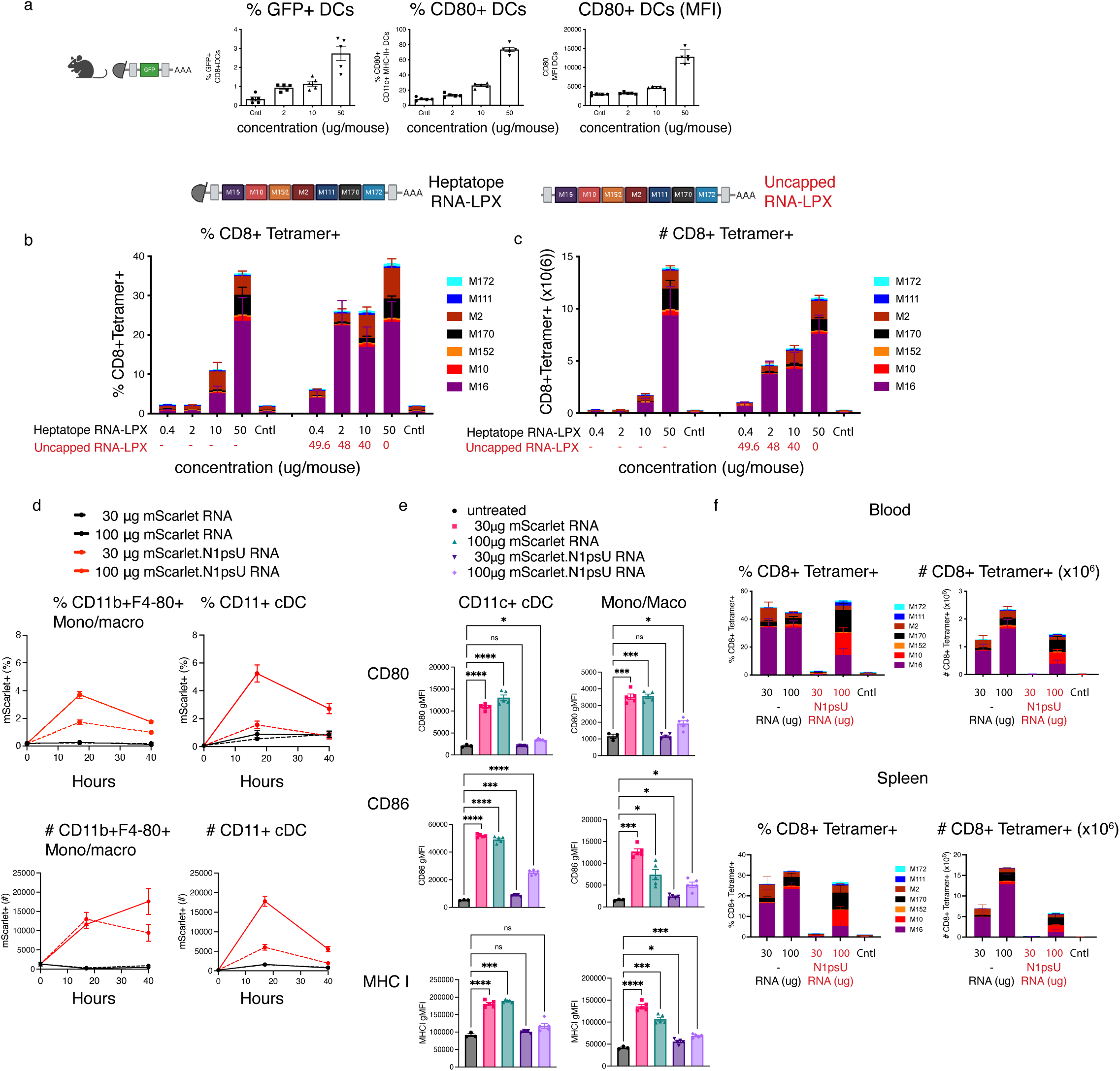
Increasing antigen expression can mitigate immunodominance hierarchies. a, C57BL/6 mice were immunized with 2, 10 or 50µg of RNA-LPX vaccine encoding GFP and DCs were evaluated in the spleen 18 h after immunization. DCs that were positive for GFP, indicating they had taken up the mRNA LPX complex were quantified by flow cytometry. The percent and MFI of the activation marker CD80 on CD11c+MHC-II+ DCs is also shown. n=5 per group. b,c C57BL/6 mice were immunized with RNA-LPX heptatope vaccine at four different concentrations, 0.4, 2, 10 or 50µg per mouse. In a second cohort, mice were immunized three times with the same four doses of heptatope RNA-LPX mixed with uncapped RNA to make a total RNA concentration of 50µg per mouse. Neoantigen-specific CD8+ T cell responses were quantified by flow cytometry in the spleen 5 days after the third immunization using pMHC-I tetramers. pMHC tetramer+ CD8+ T cells are shown as a percentage of total CD8+ T cells and the total number of neoantigen specific T cells (x106). n=5 mice. d,e C57BL/6 mice were immunized once with either 30µg or 100µg of mScarlet RNA-LPX or N1psU-modified mScarlet RNA-LPX. Mice were euthanized 17 or 40 hours after immunization and CD11c+ cDCs or a mixed population of monocytes and macrophages were examined in the spleen. n=3-5 d, Cells expressing mScarlet protein were quantified by flow cytometry. Both the percentage and total number of scarlet + cells in the spleen are shown. e, Graphs show gMFI for activation markers. f, C57BL/6 mice were immunized weekly for 4 weeks with 30µg or 100µg of either heptatope RNA-LPX or N1psU-modified hepatope RNA-LPX. Flow cytometric quantification of CD8+ T cells with pMHC tetramers was performed in the blood and spleen 4 days after the last immunization. Neoantigen-specific CD8+ T cell responses as a percentage of total CD8+ T cells and the total number of neoantigen-specific CD8+ T cells (x106) are shown (n=10 blood and n=5 spleen). Each dot represents an individual mouse. Welch’s anova. *p<0.5, **p<0.01, ***p<0.001, ****p<0.0001

To decouple the effects of innate stimulation and antigen dose on T cell responses, we co-formulated the heptatope RNA-LPX with uncapped RNA-LPX at different ratios, to keep the innate stimulation constant. Uncapped RNA-LPX was used as a “filler”, as it is not translated and does not contribute to antigen presentation but has innate adjuvanticity comparable to RNA-LPX based on serum cytokines (data not shown). At a constant high dose of innate stimulation of 50µg total, CD8+ T cell responses were detected at lower antigen doses between 2µg and 40µg in the majority of specificities (5 of 7) indicating that innate stimulation is a limiting factor for some specificities (Fig. 4b,c and Extended data Fig.4 a). The minimal dose for T cell induction still varied with the neoantigen.

Another approach to increase antigen expression is by incorporation of modified nucleotides into mRNA that reduces innate sensing, leading to enhanced mRNA stability and translation efficiency^55,56^. Administration of N1-methylpseudouridine (N1psU)-modified RNA-LPX encoding the fluorescent protein mScarlet led to enhanced and prolonged antigen expression by professional APCs compared to nucleoside-unmodified mScarlet RNA-LPX (Fig.4d). As expected, the innate immunostimulatory activity of N1psU-modified RNA-LPX was reduced, although not fully abrogated particularly at the higher concentration of 100µg where N1psU-modified RNA-LPX induced surface expression of costimulatory molecules including CD80, CD86, CCR7 and MHC molecules on professional APCs, but to a lower extent than nucleoside-unmodified RNA-LXP (Fig. 4e and Extended data Fig. 4b). Lower serum cytokines were also induced (Extended data Fig. 4c).

We next examined CD8+ T cell responses elicited by heptatope N1psU-modified RNA-LPX in comparison to nucleoside-unmodified heptatope RNA-LPX (Fig. 4f). Expectedly given the weaker innate stimulatory activity, N1psU-modified heptatope RNA-LPX vaccination induced an overall CD8+ T cell response of lesser magnitude with no T cell response detected at the lower 30µg dose, consistent with reports of reduced functionality of N1psU-modified RNA^57^. Noticeably, at the 100µg dose, the T cell hierarchy was significantly altered and the breadth increased, particularly the M10 and M170 responses. These data likely result from both enhanced antigen availability allowing for increased expansion of subdominant T cell clones, and decreased dominance of M2 and M16 responses that were more limited by innate stimulation (Fig. 4b,c).

All together, these data support the concept that antigen presentation dynamics influence immunodominance hierarchies and T cell breadth, and highlight the need for coordinated modulation of both antigen expression and innate signaling.

### T cell cross-competition drives divergent CD8+ T cell differentiation fates through immunodominance hierarchies

Since T cell cross-competition impacts the magnitude of sub-dominant T cell responses following RNA-LPX immunization, we next investigated whether this competition also alters T cell differentiation. We performed single cell RNA, ADT and TCR sequencing of purified, oligonucleotide-conjugated pMHC-I tetramer-labelled neoantigen-specific CD8+ T cells (Fig. 5a). This approach enabled us to map CD8+ T cells to their respective neoantigen specificities, yielding distributions consistent with pooled pMHC-I tetramer staining (Extended Data Fig. 5a). Clonal expansion magnitudes of the neoantigen-specificities confirmed the hierarchy observed by flow cytometry (Fig. 5b and Fig. 1b).

**Fig. 5:**
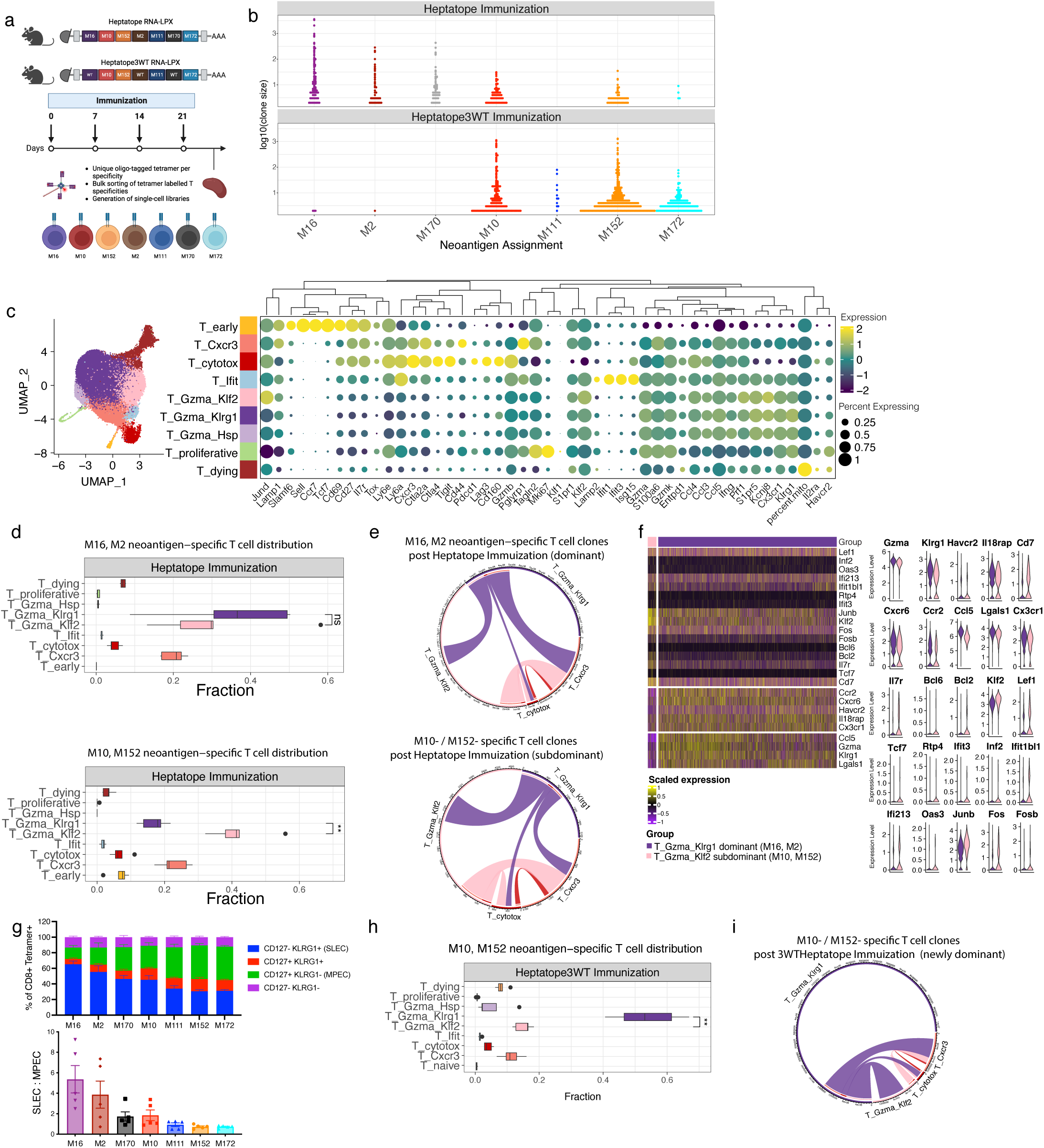
T cell cross-competition results in sub-dominant CD8+ T cells adopting a memory-precursor and stem-like phenotype. a, Experimental design for single-cell RNA, ADT, and TCR sequencing of heptatope RNA-LPX and heptatope3WT RNA-LPX vaccinated mice. Mice received four weekly vaccinations, followed by isolation and pMHC tetramer labeling of splenic CD8+ T cells for single-cell library preparation. n-5 mice per group. b, Clonal expansion beeswarm plot of neoantigen-specific T cell clones post-vaccination with heptatope RNA-LPX and heptatope3WT RNA-LPX. Each dot represents a unique clonotype based on the nucleotide sequence. Clonotypes with a single count have been excluded. c, UMAP visualization of splenic tetramer-positive T cells post-heptatope RNA-LPX and heptatope3WT RNA-LPX immunization, colored by T cell phenotype. Dotplot shows z-scored expression of selected T cell phenotype markers, dot size denotes percentage of cells with gene expression. d, Proportion of cells in each T cell phenotype relative to total cells per mouse. e, Clonal interactions among T cell phenotypes for dominant (M16, M2) and subdominant (M10, M152) neoantigen-specific T cells post-heptatope RNA-LPX immunization. Boxplot displays distribution of T cell phenotype fractions across mice, and link widths indicate shared clonotypes weighted by clone size. f, Differentially expressed genes between T_Gzma_Klrg1 M16 and M2 cells, and T_Gzma_Klf2 M10 and M152 cells, after heptatope RNA-LPX immunization, identified by pseudobulk differential expression analysis. Colors represent scaled expression values, with hierarchical clustering defining row and column order within the comparison groups. g, C57BL/6 mice were immunized three times on day 0, 7 and 14 with heptatope RNA-LPX vaccine. Flow cytometric quantification of CD127 and KLRG1 expression on neoantigen-specific CD8+ T cells from each specificity (indicated on x-axis) in the spleen on day 19 is shown. The ratio of SLEC (CD127-KLRG1+) to MPEC (CD127+KLRG1-) is shown for each neoantigen specificity. n=5 mice per group. h-i, Proportion of cells in each T cell phenotype relative to total cells per mouse and clonal interactions among T cell phenotypes for M10, M152 neoantigen-specific T cells post-heptatope3WT RNA-LPX immunization.

We identified nine distinct CD8+ T effector cell clusters, each representing a unique phenotype based on gene expression profiles (Fig. 5c, Extended Data Table 1). Similar to previous reports, we identified, (1) early T cells (T_early) expressing naive T cell markers and with minimal clonal expansion (Extended Data Fig. 5b); (2) a transitory population (T_Cxcr3) marked by *Cxcr3*, *Tigit*, *Ctla2* and the co-inhibitory molecule, *Pglyrp1*^58^; (3) cytotoxic T cells (T_cytotox) expressing *Gzmb*, *Pdcd1*, *Tigit*, *Ctla4*, and *Lag3* and, (4) interferon responsive T cells (T_Ifit) characterized by *Ifit1*, *Ifit3*, *Isg5* and *Ly6a* expression^59^. Since we enriched for only neoantigen-specific CD8+ T cells, we were able to identify two previously unidentified *Gzma*+ effector subpopulations, T_Gzma+Klrg1+ and T_Gzma+Klf2+ (Fig. 5c).

To understand how immunodominance influences fate decisions, we focused on the most abundant dominant (M16, M2) and subdominant (M10, M152) specificities (Extended Data Fig. 5a). Dominant neoantigen-specific CD8+ T cells were present in both Gzma+ subpopulations but preferentially hyper-expanded (>500 clones) within the T_Gzma+Klrg1+ cluster (Fig. 5d, Extended Data Fig. 5c, d). In contrast, subdominant neoantigen-specific CD8+ T cells were significantly enriched and expanded in the T_Gzma+Klf2+ cluster. Supporting this bifurcation in differentiation, clonal interaction analysis for the dominant M16 and M2 CD8+ T cell clones suggested equal clonal expansion into the T_Cxcr3+, T_cytotox, T_Gzma+Klrg1+ and T_Gzma+Klf2+ clusters, while subdominant M10 and M152 CD8+ T cell clonal interactions suggested a primary clonal expansion into T_Gzma+Klf2+ (Fig. 5e).

We further characterized the phenotypic difference between the T_Gzma+Klrg1+ cells of the dominant neoantigen specificities and T_Gzma+Klf2+ cells of subdominant neoantigen specificities using differential expression analysis. Dominant T_Gzma+Klrg1+ cells expressed markers associated with terminal differentiation and cytotoxicity such as *Gzma*, *Klrg1*, *Havcr2*, *Il18rap*, *Cd7*^60^, as well as *Cxcr6*, *Ccr2*, *Ccl5*, and *Lgals1*^61–64^ (Fig. 5f, Extended Data Table 2). These cells also showed increased *Cx3cr1* expression, typical of highly functional, terminally differentiated CD8+ T cells^60^. In contrast, subdominant T_Gzma+Klf2+ cells expressed markers indicative of memory and stem-like features, including *Il7r*, *Bcl6*, *Bcl2*, *Klf2*, and *Klf3*, and *Lef1*^65^, as well as *Tcf7*^66,67^, albeit restricted to a few cells consistent with our previous observations of a progressive reduction following repeated immunization^68^. In addition, interferon-stimulated genes (*Rtp4*, *Ifit3*, *Inf2*, *Ifit1bl1*, *Ifi213*, *Oas3*) were significantly upregulated, as has previously been observed in TCF1+ progenitor cells^69^, as well as AP-1 transcription factors, including *Jun* and *Fos* family members. These transcriptional features were supported at the protein level by higher IL7R (CD127) and lower KLRG1 expression among subdominant neoantigen-specific CD8+ T cells, consistent with a memory precursor effector cell (MPEC) phenotype (Fig. 5g).

To assess whether differentiation patterns were determined by intrinsic features of individual neoantigens or were a consequence of the immunodominance hierarchy, we used the heptatope3WT RNA-LPX vaccine in which the stronger neoantigens were replaced with their non-immunogenic wildtype counterparts. Under these conditions, previously subdominant neoantigen-specific T cell responses, such as M10 and M152 became enriched in the effector T_Gzma+Klrg1+ cluster demonstrating that differentiation is driven by the immunodominance hierarchy rather than neoantigen specificity (Fig. 5h, Extended Data Fig. 5c). Clonal interaction analysis reinforced this shift showing minimal representation of these newly dominant clones in the T_Gzma+Klf2+ cluster (Fig. 5i).

Taken together, these findings demonstrate that CD8+ T cell differentiation states are not fixed properties of individual neoantigens but are dynamically shaped by their position within the immunodominance hierarchy and as a result of T cell cross-competition, subdominant T cell responses are preferentially biased toward a fate with memory and stem-like features. Our results provide mechanistic insights into how inter-specificity T cell dynamics modulate both the magnitude and phenotype of vaccine-induced T cell responses, insights that may inform rational approaches for improving T cell breadth and durability in cancer immunotherapy.

## Discussion

Immunodominance is a pervasive phenomenon that plays a crucial role in shaping T cell responses, yet the underlying mechanisms remain poorly defined and likely vary depending on the T cell lineage - CD4+ versus CD8+ - which differ in key properties such as expansion potential and cytokine dependence^3,4^, as well as the immunological context such as infection, cancer or vaccination, each of which engage distinct priming environments. Consequently, it is not surprising that different determinants of immunodominance predominate across experimental models and platforms. Building on this, our study investigates how immunodominance emerges in the setting of mRNA-based vaccination and reveals that T cell cross-competition profoundly modulates the magnitude and breadth of CD8+ T cell responses while also influencing their differentiation phenotype.

In this study, we investigated how CD8+ T cell hierarchies emerge following RNA-LPX vaccination and how these hierarchies influence T cell phenotype. Using seven naturally occurring cancer neoantigens to model clinically relevant multi-neoantigen vaccine approaches, we found that immunodominance shapes CD8+ T cell responses, while leaving CD4+ T cell responses largely unaffected. Mechanistically, CD8+ T cell cross-competition between different neoantigen specificities at the time of exponential T cell expansion was a key driver of this phenomenon. We also identified the stability of pMHC-I complexes as a determinant influencing the dominance hierarchy, a finding consistent with previous reports in tumor-bearing mice^53^ , infectious disease models^10,70,71^, and other vaccine modalities^72^, suggesting that competition for limited antigen presentation is a conserved driver of immunodominance across platforms. Although pMHC-I stability broadly correlated with the immunodominance hierarchy and experimentally modulating pMHC-I stability reshaped the T cell hierarchy, the thermal denaturation stability measurements were not an exact match across all neoantigens, suggesting that additional factors may also contribute to the observed hierarchies^73–76^. For example, antigen processing and TAP transport efficiency may vary and affect the level of antigen presentation^77^.

Our results support a model in which neoantigens with greater pMHC-I stability enable prolonged T cell engagement with APCs, affording these T cells greater opportunity for expansion and competitive advantage. Given that antigen expression and presentation is relatively short lived after RNA-LPX immunization peaking within 24 hours^36^, neoantigens with more stable pMHC-I complexes may disproportionately benefit in this window. Experimental modulation of antigen availability, either through increasing the number of professional APCs expressing the neoantigens by augmenting the vaccine dose or enhancing the level and duration of antigen expression by using a nucleotide-modified mRNA enabled broader responses, strongly implicating antigen presentation as a limiting factor. However it is important to point out that nucleotide-modified mRNA also significantly reduced innate stimulation to the point of negatively impacting the overall magnitude and functionality of the T cell response (Fig. 4 and ^57^). Therefore, a careful modulation of both antigen expression and innate stimulation is required for induction of an optimal response. Prior work on T cell competition indicates that CD8+ T cells specific for different epitopes can interfere with another’s activation when antigens are presented on the same APC or in conditions of high CD8+ T cell to APC ratios^78,79^. Relatedly, it has been shown that inter-T cell competition among T cell clones targeting the same epitope can be overcome when APCs present different epitopes separately^80,81^. Based on these observations, another approach that may limit cross-competition is dose fractionation where vaccine doses are injected across multiple sites, though systemically delivered RNA-LPX vaccines overcome this problem.

We further found that immunodominance impacted the differentiation state of neoantigen-specific CD8+ T cells. We took advantage of DNA barcoding of all seven T cell specificities encoded in the heptatope RNA-LPX combined with single cell sequencing to simultaneously track clonal expansion and gene expression within each specificity^82^. Our findings revealed a divergence in differentiation where dominant CD8+ T cell responses were driven toward terminal differentiation, while subdominant CD8+ T cell responses adopted memory-precursor and stem-like features, a gene expression profile consistent with less functional potential, extending previous findings in viral and tumor models^53,83^. Notably, these phenotypes were not fixed. We found that sub-dominant T cell responses possessed the intrinsic capacity to adopt the ‘dominant’ T_Gzma+Klrg1+ terminal effector molecular phenotype when dominant competitors were absent, consistent with a dynamic definition of dominant and subdominant that is dictated entirely by the attributes of the other concurrent T cell responses. While we did not directly assess in vivo cytotoxic function or anti-tumor efficacy of dominant and subdominant specificities, the finding of divergent T cell fates and transcriptional profiles consistent with altered functional potential provides a mechanistic basis for future functional studies. It has previously been reported that the onset of IFNγ production within epitope-specific CD8+ T cell populations correlates with their position within the immunodominance hierarchy^84^. Collectively, our data underscores that rather than being a rigid determinant of T cell fate, immunodominance hierarchies dynamically influence the trajectory of T cell responses.

These findings carry important implications for therapeutic vaccine design for cancer and infectious disease. Achieving broad CD8+ T cell responses against multiple antigens is likely critical to overcome tumor heterogeneity and immune escape^85^, a notion supported by clinical data suggesting that increased T cell breadth is associated with better outcomes^86–91^. Our data highlight how features like pMHC-I stability, innate stimulation and T cell cross-competition can shape this breadth and suggest that these parameters may inform rational antigen selection. For example, choosing antigens with comparable predicted pMHC-I stability might reduce competition and favor co-dominant responses. Additionally, future studies may investigate whether adjusting antigen presentation kinetics or leveraging innate adjuvants could fine-tune T cell differentiation to favor memory or effector profiles.

These insights underscore the plasticity of T cell fate and highlight the importance of considering competitive dynamics in multi-neoantigen vaccine design. While our findings were derived in the context of RNA-LPX vaccines, the mechanisms described are likely applicable across diverse vaccine platforms, and offer a framework for optimizing vaccine-induced immunity in both cancer and infectious disease settings.

## Materials and Methods

### Mice

C57BL/6 mice (stock 000664) were purchased from Charles River. Age-matched (6–10 weeks) female animals were used throughout all experiments. Mice were maintained in accordance with the Guide for the Care and Use of Laboratory Animals (National Research Council, 2011). Genentech is an American Association of Laboratory Animal Care-accredited facility and all animal activities in this research study were conducted under protocols approved by the Genentech IACUC. Mice were maintained in a specific-pathogen-free facility, in individually ventilated cages within animal rooms maintained on a 14-h/10-h, light/dark cycle. Animal rooms were temperature- and humidity-controlled, at 68–79 °F and 30–70%, respectively, with 10 to 15 room air exchanges per hour. Female mice (aged 6–10 weeks) that seemed healthy and free from obvious abnormalities were used for the study.

### Vaccination

For RNA-LPX vaccination, RNAs were synthesized by Genentech and were formulated with liposomes consisting of DOTMA and DOPE at a charge ratio (+):(−) of 1.3:2, yielding negatively charged RNA-LPX^36^. For some experiments, RNA was synthesized using N1-methylpseudouridine instead of uridine (N1psU RNA-LPX) and RNA was purified with cellulose treatment to remove dsRNA contaminants. Unless otherwise indicated, mice were intravenously immunized with 50µg of RNA-LPX weekly for three weeks, and were euthanized four to seven days after the final immunization.

### Spleen and blood preparation

Spleens were collected in cold PBS and single-cell suspensions were generated by mashing the spleen tissue through a 70-μm cell strainer (BD Falcon) in Hank’s-based Cell Dissociation Buffer (Gibco) supplemented with Liberase (Roche) and DNase I (ThermoFisher). Red blood cells were lysed with ACK lysis buffer (Gibco). For immune cell analysis, retro-orbital bleeding under isoflurane anesthesia was used to collect peripheral blood samples on day 5 following each immunization. Whole blood was stored in lithium heparin microtainer tubes (BD) and red blood cells were lysed with ACK lysis buffer (Gibco). In some experiments, mice were bled for cytokine analysis 4–6 h after the first immunization. Blood was stored in gel-separator with clot activator polypropylene tubes (Sarstedt) and incubated for 15 min at room temperature, after which the coagulated blood samples were centrifuged at 2,300g for 5 min. Clear serum was transferred to new tubes and stored at −80 °C for downstream assays.

### Flow cytometry

Single-cell suspensions were incubated in FACS buffer (PBS supplemented with 0.5% BSA and 0.05% sodium azide) containing anti-mouse CD16/CD32 (Mouse Fc Block, BD) for 10 min before staining with the indicated antibodies. Cells were stained on ice for extracellular markers for 20 min. In some experiments, cells were further stained for intracellular markers for 30 min. Cells were filtered using 30–40-μm filter plates (PALL). Samples were acquired with BD FACSDiva software v.8.0 on a BD FACSymphony (BD) and analyzed with Flow Jo v.10.7.1 (TreeStar). Dead cells and cell aggregates were excluded from analyses by LIVE/DEAD Fixable Near-IR (Dead Cell Stain Kit for 633- or 635-nm excitation, Invitrogen) staining and forward scatter area (FSC-A)/forward scatter height (FSC-H) characteristics.

To identify neoantigen-specific T cells, cells were stained with fluorochrome-conjugated pMHC-I tetramers (Genentech) for 20 min at room temperature, followed by cell surface marker staining as described above. For intracellular cytokine detection, spleen cells from immunized mice were stimulated with neoantigen peptides at 37C for 4 hrs. A mixture of 1X Golgi plug and 1X Golgi stop (BD Biosciences) and recombinant IL-2 (Sigma) was added for the duration of the 4 h stimulation. Intracellular Fixation & Permeabilization Buffer Set (eBioscience) was used for intracellular staining of IFNγ production according to manufacturer’s instructions. Antigen presenting cells were identified by flow cytometry using the following gating strategies: DCs (Live+, CD45+, Dump-, SiglecH-, F4-80-, CD11c+, MHC-II+ and a mixed population of monocytes and macrophages (Live+, CD45+, Dump-, SiglecH-, CD11b+, F4-80+).

Cells were purified by fluorescence-activated cell sorting (FACS) on a Becton Dickinson FACSAria Fusion cell sorter equipped with four lasers (405 nm, 488 nm, 561 nm and 638 nm). A 70-μm nozzle running at 70 psi and 90 kHz was used as the setup for each sort session. Before gating on fluorescence, single cells were gated using forward scatter (FSC-A) and side scatter (SSC-A) (for intact cells) and SSC-W/SSC-H and FSC-W/FSC-H (to ensure that only singlets were sorted).

### IFNγ ELISpot

5 × 10^5^ splenocytes per well were cultured overnight at 37°C in complete RPMI 1640 media (10% FBS, 1% penicillin/streptomycin, 1% Hepes, 1% GlutaMAX, and 1% sodium pyruvate) in pre-coated 96-well mouse IFNγ ELISpot plates (R&D Systems). For stimulation, 27mer synthetic long peptides containing the neoantigen were added (10 µg/ml in DMSO stock). For analysis, IFNγ spots were developed according to the manufacturer’s protocol and counted using an automatic ELISpot reader (AID). Peptide synthesis was performed by Anaspec (75% purity).

### Luminex cytokine and chemokine assay

Serum concentrations of murine cytokines and chemokines were determined using a bead-based, Cytokine & Chemokine Convenience 26-Plex Mouse ProcartaPlex multiplex immunoassay (ThermoFisher Scientific) according to the manufacturer’s instructions. Values below the lower limit of quantification were set to zero. Luminex data were collected on Flex-Map 3D v.3.2 (Luminex Corporation) and analyzed with Bio-Plex Manager 6.1.1 and Microsoft Excel v.16.16.27.

### Differential scanning fluorimetry (DSF) assay

H-2Kb and H-2Db peptide complexes were made using UV-mediated ligand exchange as described previously^92,93^, formulated in 25mM TRIS pH 8.0, 150mM NaCl, 4mM EDTA, 5% ethylene glycol. After exposure to UV light, the peptide exchange reaction was allowed to proceed for 18 hrs at room temperature. Melting temperatures were determined using differential scanning fluorimetry which was performed on a Bio-Rad CFX96RT C1000 Touch qPCR machine monitoring fluorescence at Ex/Em = 587/607 nm. 24 ul of 50 µg/ml solutions of H-2kb

or H-2Db peptide complexes were added to a Bio-Rad hard-shell 96-well PCR plate to wells containing 1ul of 25X SYPRO Orange and sealed with a Bio-Rad Microseal “C” sealing film. Thermostability measurements were acquired over a temperature gradient of 20℃ to 100℃ at a rate of 0.2℃ per 10 seconds. Data analysis was performed on CFX Manager software and Tm was calculated using the negative first derivative of RFU values over temperature. Melting curves were normalized to the minimum and maximum fluorescence values.

To evaluate the M16 neoantigen and the mutated M16-9L variant, H2Db heavy chain, β2m and the corresponding peptide were refolded together in refolding buffer (0.4 M arginine, 0.1 M Tris, GSH:GSSG = 5 mM:0.5 mM, 2 mM EDTA, 0.5 mM PMSF, pH 8.0) for 1 week and purified as previously described. Differential scanning fluorimetry was performed using QuantStudio 12 Flex (Thermo Fisher) with Protein Thermal Shift TM dye (Thermo #4461146). Briefly, 12.5 µl of protein in PBS with a concentration 0.3 - 0.5 mg/ml was mixed with 2.5 µl of 8X dye solution (1:1000 dilution in PBS buffer) and 5 µl reaction buffer. The melting curves were measured from 25 °C to 99 °C with a temperature scan rate of 0.05 °C/s, and three duplicates were performed. The melting temperature (Tm) is calculated from a plot of d(fluorescence)/dT to the experimental raw curve.

### Peptide-induced H2-b stabilization assay

Tap1-knockout EL4 cell lines were generated as previously described^94^. A sub-clone was further engineered to overexpress H2-Db and H2-Kb to increase the dynamic range of the stabilization assay. Briefly, lentivirus encoding H2-b alleles were generated (pLKO_Ef1) in 293 cells. Supernatants containing the virus were concentrated using Lenti-X (Takara Bio). Tap1-knockout EL4 cells were infected with lentivirus followed by puromycin selection. H2-b overexpression was confirmed by H2-b surface expression relative to mock-transduced cells.

For testing the relative affinities of H2-b peptides, cells were incubated overnight at 26°C, 5% CO2 to promote accumulation of peptide-free H2-b surface molecules. 2 × 10^5^ cells per well were seeded into 96-well U-bottom plates containing 10 µg/ml β2-microglobulin (WuXi Biologics) with 9mer neoantigen peptides (Anaspec) over a range of 10-fold dilutions in a total volume of 100 µl of complete RPMI 1640 culture medium. Cells were cultured for 4 h at 26°C, 5% CO2 and a further 2h at 37°C, 5% CO2 to promote denaturation of empty H2-b complexes. The surface expression of H2-b, quantified as geometric median fluorescence intensity, was determined by flow cytometry using PE conjugated anti-mouse H-2Kb antibody (Clone AF6-88.5.5.3, BD Biosciences) and H2-Db (clone KH95, BioLegend). After staining, cells were fixed in 1% PFA. OVA_257-264_ (H2-Kb) and LCMV_gp33-41_ (H2-Db) peptides were purchased from MBL and used as positive controls. Isotype controls were used to subtract background staining. Data are presented as percent of maximum response at 10μg/ml.

### Naïve precursor cell isolation and quantification

The total pre-immune precursor T cell repertoire for each of the neoantigen specificities was quantified as previously described using a combination of APC labelled pMHC-I tetramer staining and magnetic particle-based cell enrichment^95,96^. Briefly, all secondary lymphoid organs from a non-immunized mouse were removed and single cell suspensions were prepared. Following pMHC-I tetramer staining in Fc block for 1 hr at RT, anti-APC microbeads (Miltenyi Biotec 130-090-855) were used to enrich precursor cells on LS columns (Miltenyi Biotec 130-042-401). The bound cells were eluted and the precursor fraction of CD8+ T cells was quantified by flow cytometry as described^97^. Beads (Spherotech ACFP-70-5 (7.9um particles)) were used to quantify the total number of cells in the bound fraction of the enriched sample.

### Single-cell RNA Sequencing

Following immunization, splenic CD8+ T cells were enriched by depletion of B and CD4+ T cells using magnetic beads (Stemcell Technologies; Robosep). The remaining CD8+ T cells were stained with a mixture of the seven MHC-I/neoepitope tetramers that had been conjugated with streptavidin-PE containing an oligonucleotide barcode label (TotalSeq^TM^-C, Biolegend) so that each of the 7 neoantigen specificities could be identified by a unique barcode. All MHC-I/neoepitope tetramer positive cells were FACS sorted, and single cells were partitioned into Gel Bead-in-Emulsions (GEMs) using the Chromium single cell 5’ chip K on a Chromium controller (10X Genomics), followed by cell lysis and barcoded reverse transcription of mRNA. Paired single cell 5’ gene expression, antigen-binding (feature barcode) and VDJ sequence libraries were generated according to manufacturer’s instructions using the following kits: Chromium Next GEM Single Cell 5’ Kit v2 (1000263), Chromium Single Cell Mouse TCR Amplification Kit (1000254), Library Construction Kit (1000190), 5’ Feature Barcode Kit (1000256), Dual Index Kit TN Set A (1000250) and Dual Index Kit TT Set A (1000215) (10X Genomics). Gene expression libraries were sequenced on an Illumina NovaSeq 6000 instrument at read lengths of 28x90+10+10 and a sequencing depth of 20k per cell. scTCRseq and ADT libraries were sequenced on an Illumina NovaSeq 6000 and Illumina HiSeq 4000 instrument, respectively. FASTQ files of gene expression, feature barcode and TCR libraries were generated and analyzed with CellRanger (v7.1.0) using the GRCm38 mouse reference genome, barcode feature reference and VDJ GRCm38 5.0.0 for alignment, respectively.

### scRNA-seq processing and analysis

The CellRanger output filtered barcode matrices were demultiplexed using DemuxEM v1.12.0^98^ and further processed together with the filtered feature expression matrices with Seurat v.5.0.1^99^. Briefly, library barcode antibodies for cell hashing were demultiplexed and assigned to their respective neoantigen specificities. Rare cases in which cells from the same clonotype were assigned to different neoantigen specificities were resolved by correcting the assignment to the majority vote. Individual libraries were merged into a Seurat object and filtered for neoantigen-specific singlets with over 300 features and less than 5% mitochondrial transcripts. Feature counts were normalized per cell, scaled by 10,000, and log-transformed. T-cell receptor genes were excluded from further analysis. The top 2000 highly variable features were identified using variance stabilizing transformation, followed by data scaling, principal component analysis, and Uniform Manifold Approximation and Projection (UMAP) dimensionality reduction with 30 principal components. Clustering was performed using shared nearest neighbors and graph-based methods. Clusters suspected of cell contaminants with expression of *Cd74*, *Xist*, *Hba-a1*, *Hba-a2* were filtered and data reprocessed as above. T cell phenotypes were characterized at a clustering resolution of 0.3 by a combination of unbiased cluster marker analysis and supervised analysis of T cell phenotype and function gene expression.

Differential gene expression analysis was performed using a pseudobulk differential expression analysis approach utilizing limma v3.58.1. In brief, the seurat count matrix was filtered to exclude non-protein-coding genes, ribosomal genes and lowly expressed genes. Counts were then aggregated by metadata groups into a pseudobulk count matrix followed by limma voom differential expression analysis using library size as a normalization factor.

Analysis of T cell phenotype fractions within a treatment group was conducted by calculating the proportion of cells relative to the total cell count in each mouse, optionally grouped by clonotype expansion (as described below). Statistical significance was assessed using the Wilcoxon Rank-Sum test, with significance levels defined as follows: *P ≤ 0.05, **P ≤ 0.01, ***P ≤ 0.001 and ****P ≤ 0.0001.

### scTCR-seq processing and analysis

The CellRanger filtered contig annotation matrices were processed using scRepertoire v2.0.4 by combining the TCR sequences, stripping the barcodes and adding the TCR information to the merged seurat object processed as described above. Clonal assignments were made using the clone nucleotidesequence (CTnt). For clonotype analysis, cells without an assigned clonotype, cells with more than one TRB and more than two TRA were removed. Clonal expansion groups were defined as single (CTnt frequency <= 1), expanded (CTnt frequency <= 500) or hyperexpanded (CTnt frequency > 500).

To compute clonal interactions and capture shared phenotypes across T cell clones, a phenotype pair matrix was computed by summing x_k_*x ^T^ over all clones, where x_k_ represents the vector of phenotype counts for clone k, and x_k_*x ^T^ represents the outer product of the vector x_k_ with its transpose x_k_ ^T^, effectively measuring the shared presence of clonotypes between pairs of phenotype clusters. The final phenotype pair matrix was visualized as ChordDiagrams using the circlize package v0.4.16.

### Statistical analyses and data presentation

Statistical analyses and graphing were performed using GraphPad Prism v.9.4.1 for Mac OS. Illustrations were created with BioRender.com. FACS analysis was performed using FlowJo v10.8.0. Unless otherwise indicated, results are expressed as mean ± s.e.m. or median values. Unpaired two-tailed Student’s t-test, one way or Welch’s anova was used as appropriate for comparison between groups. Functional avidity EC50 values were calculated using a five-parameter logistic equation where x-values are concentrations. *P ≤ 0.05, **P ≤ 0.01, ***P ≤ 0.001 and ****P ≤ 0.0001. No statistical methods were used to predetermine sample size for animal experiments.

## Data availability

Single-cell RNA, ADT, and TCR-sequencing data generated in this study will be deposited in the Gene Expression Omnibus database and accession numbers will be made available upon manuscript acceptance.

## Code availability

No new algorithms were developed as part of this study.

## Acknowledgements

We thank Mahesh Yadav and Beatrice Breart for comments and all of the members of the former Cancer Immunology Department at Genentech for support and advice.

---

**Extended Data Fig. 1.**
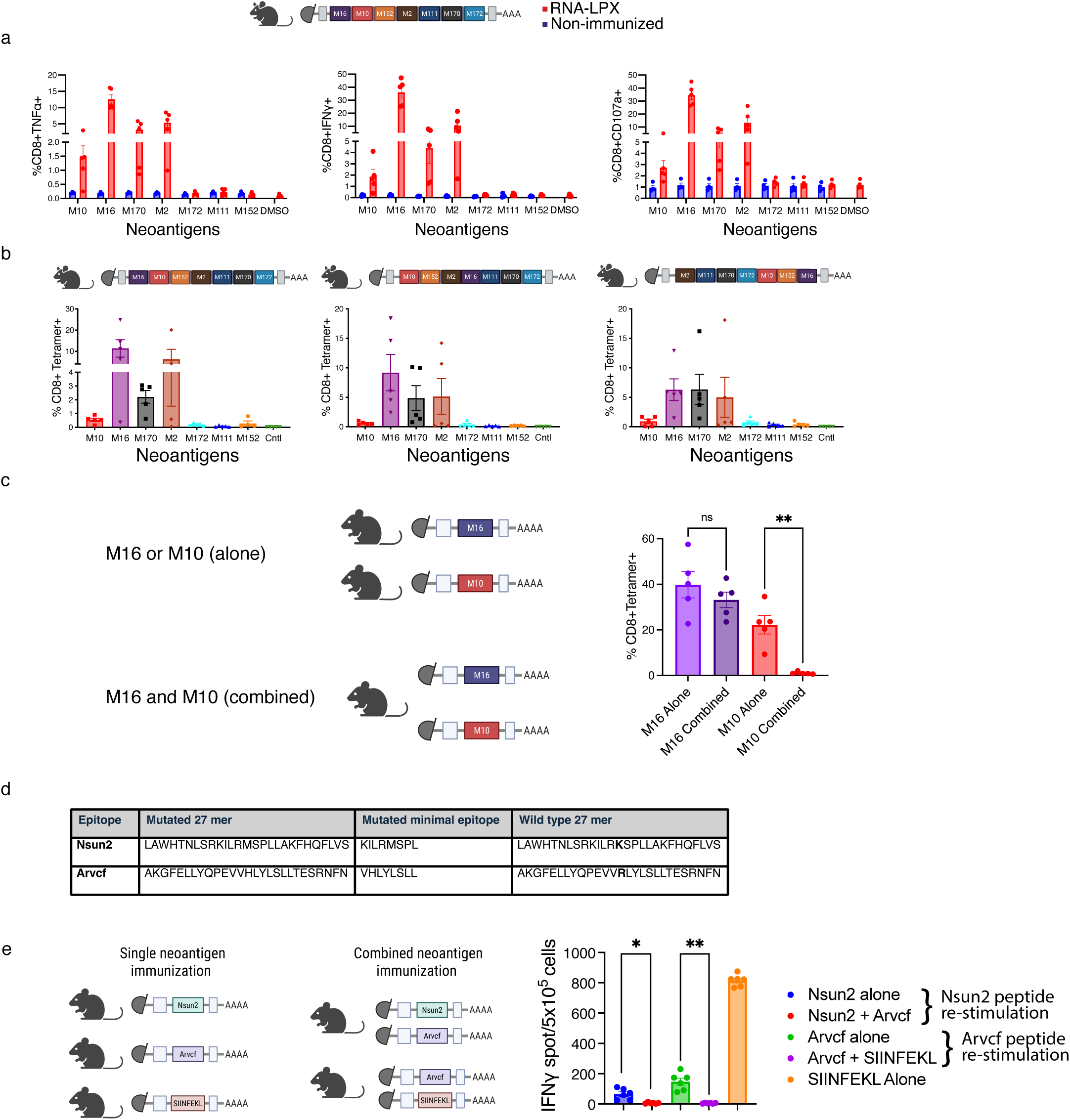
Distinct hierarchy of CD8+ T cell responses following multi-neoantigen RNA-LPX vaccination. a, C57BL/6 mice were immunized three times on days 0, 7 and 14 with heptatope RNA-LPX vaccine encoding all 7 neoantigens. Splenic T cells were harvested 5 days after the last immunization and restimulated with 27 mer peptides containing the neoantigen or non-peptide DMSO solvent controls. The magnitude of cytokine and CD107a responses were measured by flow cytometry and are shown as a percentage of total CD8+ T cells. Neoantigen specificities are indicated on the x-axis. n=3-5. b, As depicted, three different heptatope constructs were generated encoding 7 neoantigens in which the neoantigen position was varied across constructs. C57BL/6 mice were immunized three times on days 0, 7 and 14 and splenic T cells were harvested 5 days after the last immunization. Flow cytometric quantification of CD8+ T cells stained with pMHCI tetramers (specificities indicated on x-axis) was performed. n=5. c, Mice were immunized with monotope RNA-LPX vaccines encoding M16 or M10 alone or both monotype RNA-LPX vaccines were combined. Flow cytometric quantification of CD8+ T cells stained with pMHCI tetramers was performed in the blood 5 days after the third immunization. Clonal expansion was analyzed. n=5. d, Table depicts neoantigen and non-neoantigen wild type sequences of Nsun2 and Arvcf peptides identified in B16F10 tumors. e, Mice were immunized with monotope RNA-LPX vaccines encoding Nsun2, Arvcf or SIINFEKL in various combinations as depicted. Following three immunizations, splenocytes were assessed by IFNg ELISpot assay following peptide restimulation. One way anova or unpaired t-test. ns not significant, *p<0.5, **p<0.01, ***p<0.001.<0.01, ***p<0.001.

**Extended Data Fig. 2.**
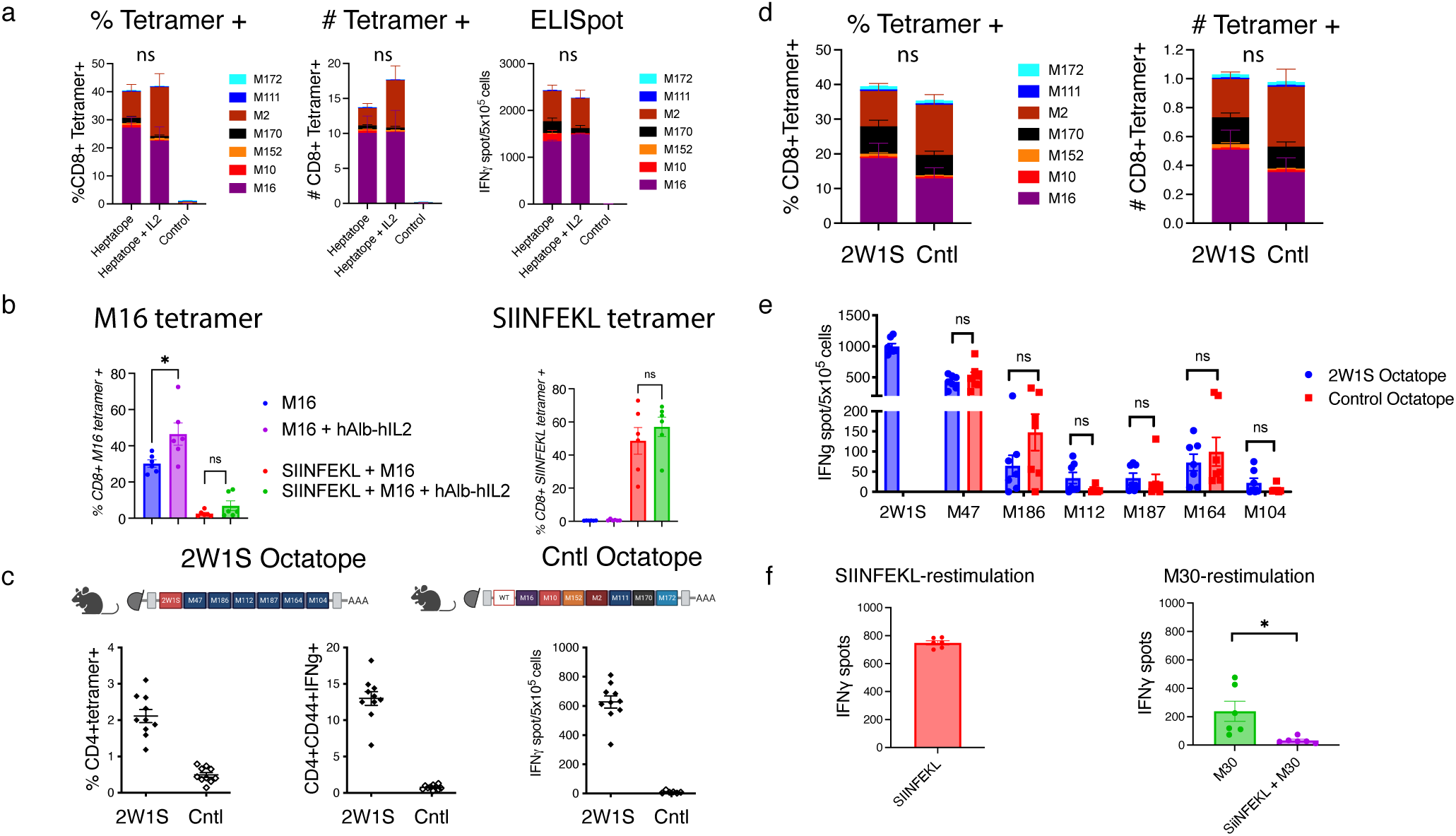
T cell competition is not mediated by the limited bioavailability of IL-2 or limitations in CD4+ T cell help. a, C57BL/6 mice were immunized with heptatope RNA-LPX in the presence and absence of exogenous IL-2 (15,000 units I.P daily). Following three immunizations, spleens were removed and neoantigen-specific T cell responses were evaluated by pMHC-I tetramer staining and IFN-gamma ELISpot. Data are shown as pMHCI tetramer+ cells as a percentage of total CD8+ T cells, total number of neoantigen-specific CD8+ T cells or IFNɣ spot count per 105 splenocytes. n=10.b Mice were immunized with monotype vaccines encoding M16 and SIINFEKL, either alone or mixed together. Mice immunized with the M16 monotype or a mixture of M16 + SIINFEKL received half-life extended human IL-2 delivered in LNP (3ug) intravenously. M16 or SIINFEKL neoantigen-specific T cell responses were evaluated in the spleen 7 days after the last immunization using pMHCI tetramer+ staining and are shown as a percentage of total CD8+ T cells. c-e, The heptatope RNA-LPX vaccine was modified to include 2W1S (“octatope”), or a non-immunogenic control antigen. Mice were immunized three times and neoantigen-specific T cell responses were evaluated in the spleen 5 days after the last immunization. c, The 2W1S CD4+ T cell response was confirmed by pMHC-II tetramer staining and production of IFNg after ex vivo restimulation of spleen cells by 2W1S peptide n=10. d, Neoantigen-specific CD8+ T cell responses were quantified in the spleen by pMHC tetramer+ staining in the presence and absence of the 2W1S CD4+ T cell responses. Data are shown as pMHCI tetramer+ cells as a percentage of total CD8+ T cells and total number of neoantigen-specific CD8+ T cells n=4-7. e, We generated a novel heptatope consisting of 6 neoantigens from the MC38 model that generate CD4+ T cell responses and a seventh CD4+ T cell antigen, 2W1S. Mice were immunized three times and neoantigen-specific CD8+ T cell responses were evaluated in the spleen 5 days later. Neoantigen-specific CD8+ T cell responses were quantified by IFNɣ ELISpot and are shown as IFNg spots per 105 splenocytes n=7. f, RNA-LPX monotype vaccines encoding either SIINFEKL or the M30 neoantigen, or both monotope vaccines combined were used to immunize mice 3 times. Neoantigen-specific T cell responses were quantified in the blood 5 days after the third immunization. n=6. Unpaired, two-tailed t-test. ns; not significant *p<0.5.

**Extended Data Fig. 3.**
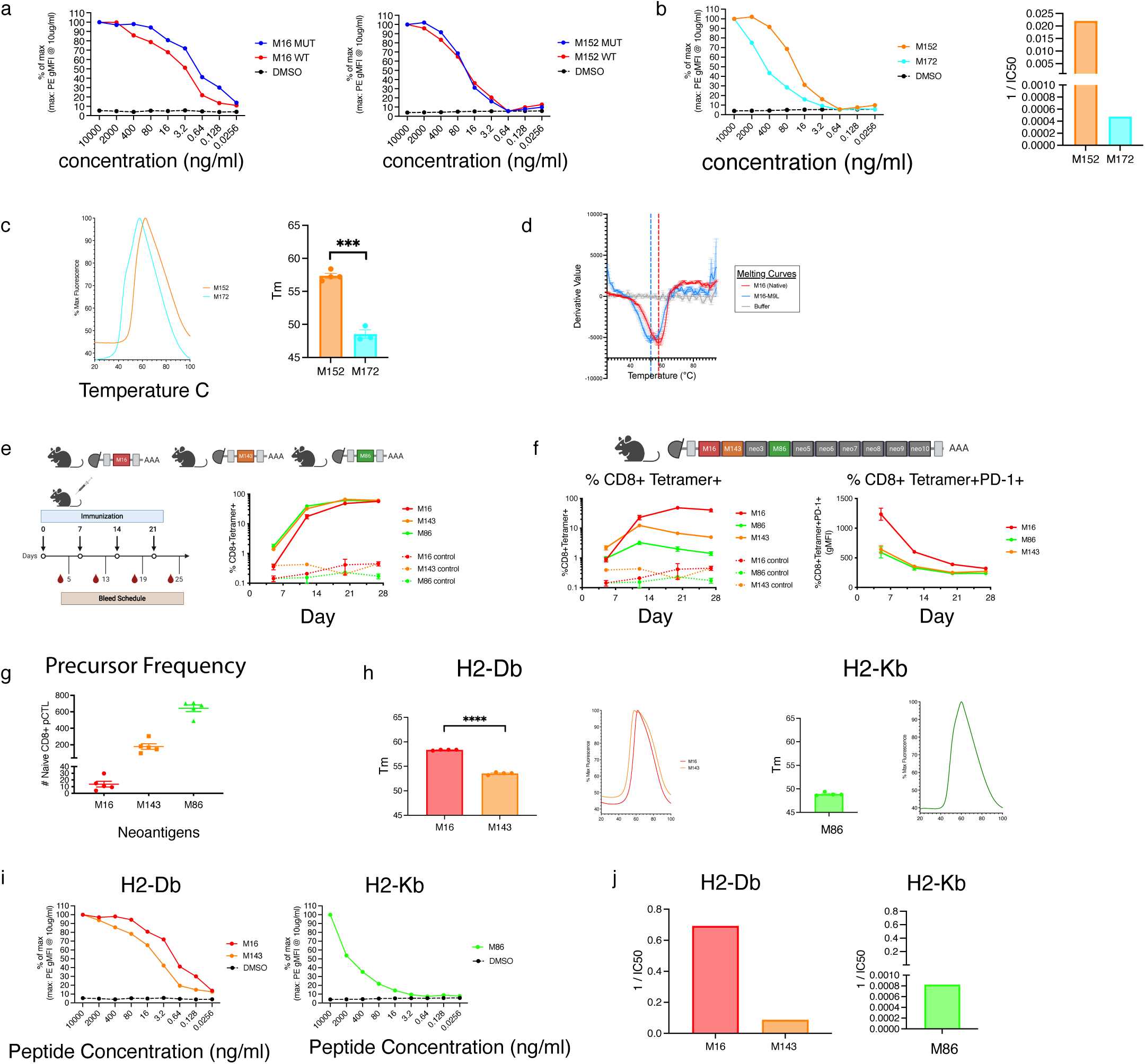
Neoantigen-MHC class I stability partially determines immunodominance. a, pMHC affinity for each neoantigen was assessed by flow cytometric quantification of H2-Db or H2-Kb stabilization for both wild type (WT) or neoantigen mutant (MUT) peptides on TAP-deficient EL4 cells. Graph shows percent of maximum response.b, pMHC affinity for each neoantigen was assessed by flow cytometric quantification of H2-Kb stabilization of TAP-deficient EL4 cells. Percent of maximum response over a range of neoantigen concentrations is shown and the graph represents the calculated 1/EC50 for each specificity. A representative graph of three independent experiments is shown. c, pMHC complex stability (Koff) for each H2-Kb neoantigen was measured by differential scanning fluorimetry following thermal denaturation of soluble pMHC-I complexes. Melting curves were normalized to the minimum and maximum fluorescence values and Tm was calculated using the negative first derivative of RFU values over temperature. n=3.d, Differential scanning fluorimetry analysis of H-2Db/M16 (red) and H2-Db/M16-9L (blue) and buffer control (grey). Curves show the derivative of fluorescence over temperature, with dotted lines marking the melting temperature (T_m_) on the x-axis. Each trace represents triplicate measurements. e, C57BL/6 mice were immunized four times on days 0, 7, 14 and 21 with either RNA-LPX monotope vaccines encoding a single neoantigen, decatope RNA-LPX vaccine encoding 10 neoantigens or RNA-LPX encoding F-luciferase was used as a control. Experimental design and flow cytometric quantification of CD8+ T cells with pMHCI-tetramer staining was performed in the blood 5 days after each immunization. Neoantigen specific CD8+ T cell responses as a percentage of total CD8+ T cells n=5. f, Neoantigen specific CD8+ T cell responses as a percentage of total CD8+ T cells is shown. Graph shows PD1 (gMFI) expression on neoantigen-specific CD8+ T cells=5. g, Naïve precursor CD8+ T cells (pCTL) were enriched from non-immunized mice and quantified. Graph shows the total number of pCTL for each specificity. Each dot represents an individual mouse n=5. h, pMHC complex stability (Koff) for each neoantigen was measured by differential scanning fluorimetry following thermal denaturation of soluble pMHC-I complexes. Melting curves were normalized to the minimum and maximum fluorescence values and Tm was calculated using the negative first derivative of RFU values over temperature. n=4 independent studies. i,j pMHC affinity for each neoantigen was assessed by flow cytometric quantification of H2-Db stabilization of TAP-deficient EL4 cells. Graph shows percent of maximum response and graph shows 1/EC50. ns not significant, *p<0.5, **p<0.01, ***p<0.001.<0.01, ***p<0.001.

**Extended Data Fig. 4.**
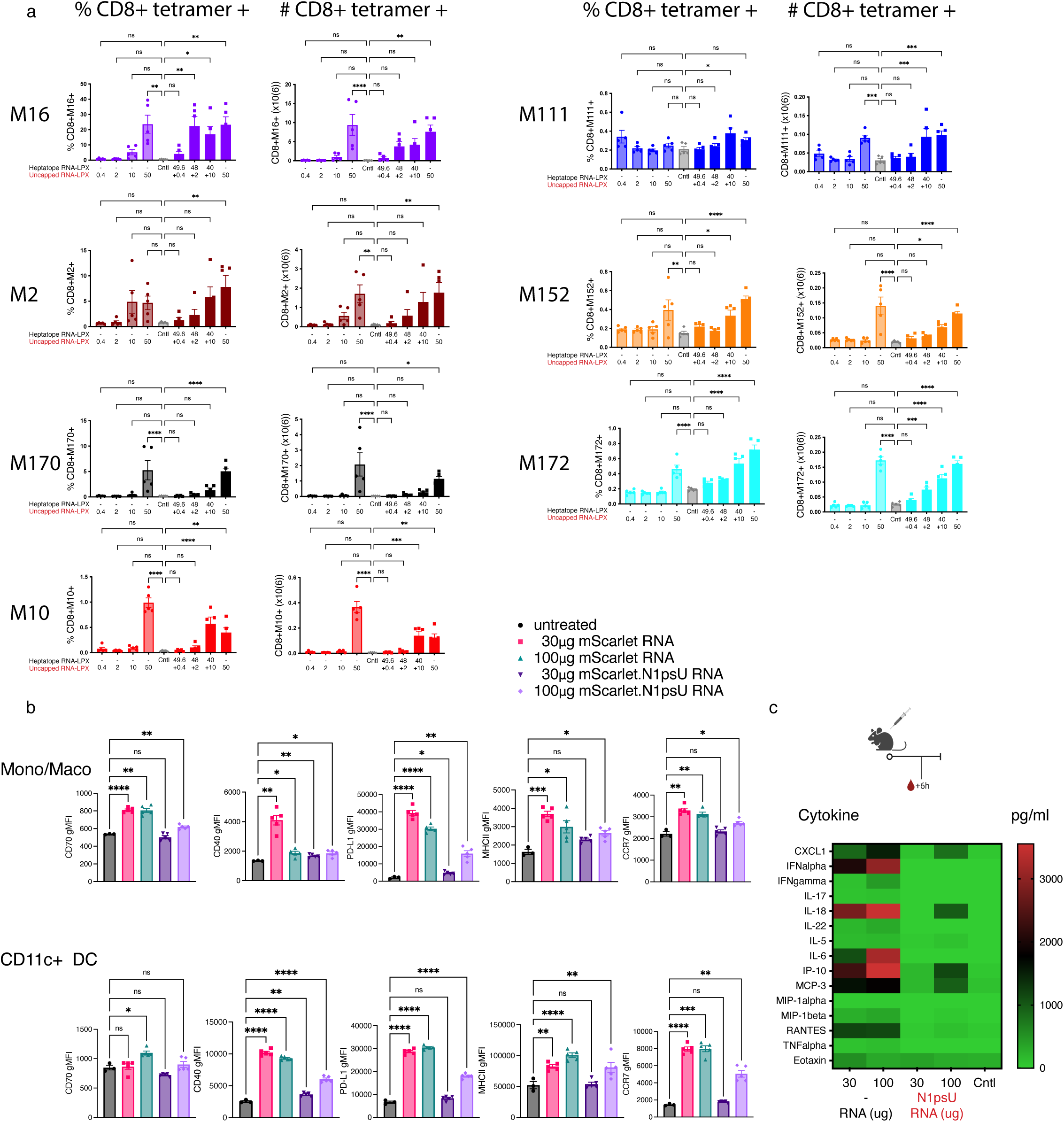
a, C57BL/6 mice were immunized with RNA-LPX heptatope vaccine at four different concentrations, 0.4, 2, 10 or 50µg per mouse. In a second cohort, mice were immunized three times with the same four doses of heptatope RNA-LPX mixed with uncapped RNA to make a total RNA concentration of 50µg per mouse. Neoantigen-specific CD8+ T cell responses were quantified by flow cytometry in the spleen 5 days after the third immunization using pMHC-I tetramers. pMHC tetramer+ CD8+ T cells are shown as a percentage of total CD8+ T cells and the total number of neoantigen specific T cells (x106). n=5 mice.b, C57BL/6 mice were immunized once with either 30µg or 100µg of mScarlet RNA-LPX or N1psU-modified mScarlet RNA-LPX. CD11c+ DCs or a mixed population of monocytes and macrophages were examined in the spleen. Graphs show gMFI for activation markers.n=3-5. c, Mice were immunized once with either 30µg or 100µg of heptatope RNA-LPX or N1psU-modified hepatope RNA-LPX. Six hours later, mice were bled and serum cytokines and chemokines were measured by luminex. Heatmap is based on the median expression of n=10 individual mice. Welch’s anova. *p<0.5, **p<0.01, ***p<0.001, ****p<0.0001

**Extended Data Fig. 5.**
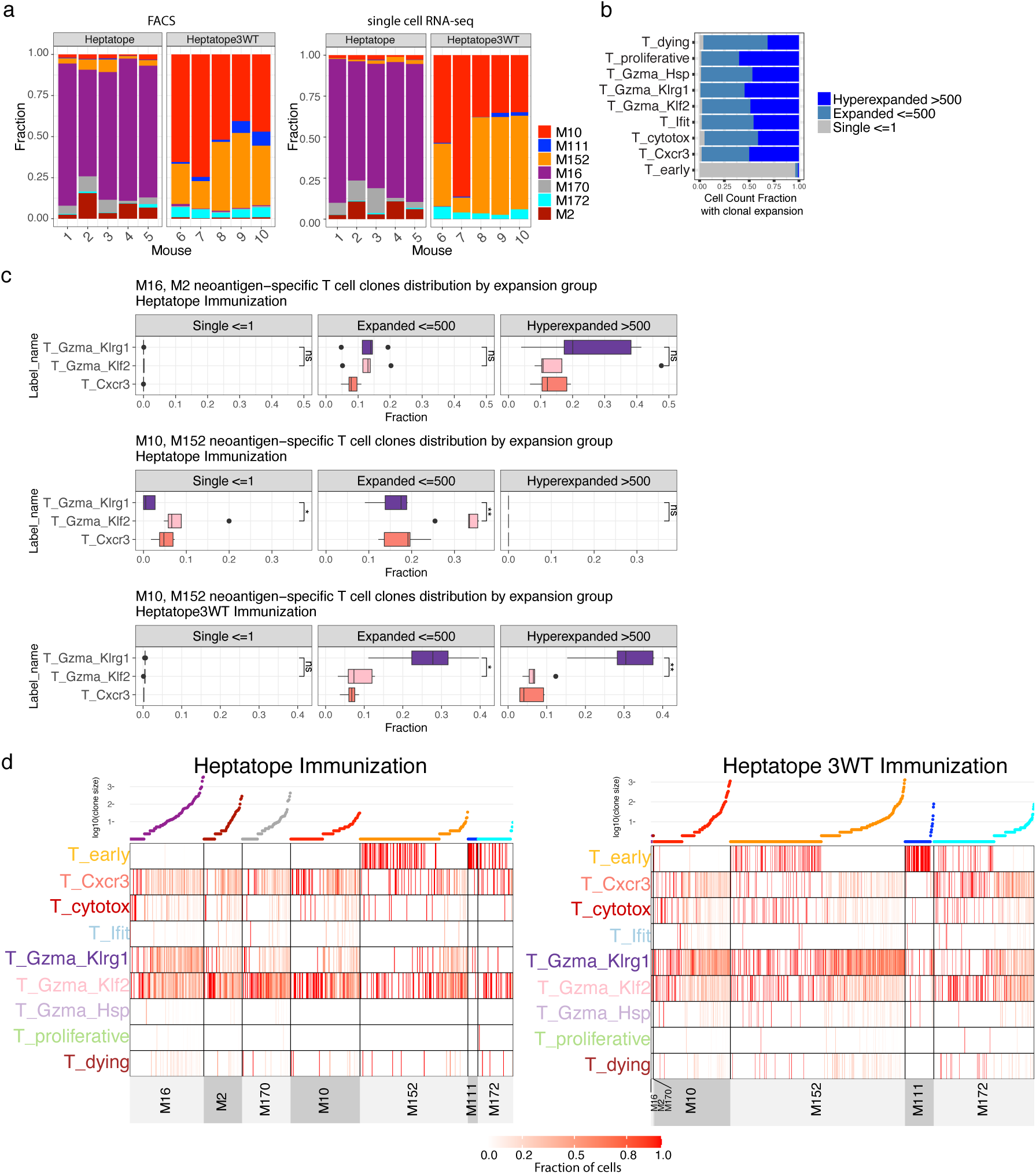
Clonal expansion tracks with preferential differentiation of subdominant CD8+ T cells into a memory-precursor and progenitor-like phenotype. a, Cell count fraction of neoantigen-specific tetramer-positive T cells after heptatope and heptatope3WT RNA-LPX immunization, as quantified by FACS and analyzed through single-cell RNA/ADT/TCR sequencing. b, Distribution of clonal expansion across T cell phenotypes, shown as the fraction of cells with an assigned clonotype expansion: single (≤ 1 cell with clonotype), expanded (≤ 500 cells with a clonotype), hyperexpanded (> 500 cells with a clonotype). c, Clonal expansion of cells with a specific T cell phenotype and neoantigen-specificity represented as the fraction of cells in each clonotype expansion group, T cell phenotype, and neoantigen-specificity relative to the total number of cells with a given neoantigen-specificity and T cell phenotype in a mouse. d, Distribution of neoantigen-specific clonotypes across T cell phenotypes as heatmap with columns representing each specific clonotype of a neoantigen-specificity and rows representing the T cell phenotypes. Colors indicate the fraction of cells of a specific clonotype in a T cell phenotype relative to all cells of that clonotype and neoantigen-specificity. Clonotypes are ordered by increasing clone size, with a scatterplot above the heatmap illustrating the log10 clone size of each clonotype.

## References

1. Yewdell, J. W. & Bennink, J. R. IMMUNODOMINANCE IN MAJOR HISTOCOMPATIBILITY COMPLEX CLASS I–RESTRICTED T LYMPHOCYTE RESPONSES1. Annu Rev Immunol 17, 51–88 (1999).

2. Kavazović, I., Polić, B. & Wensveen, F. M. Cheating the Hunger Games; Mechanisms Controlling Clonal Diversity of CD8 Effector and Memory Populations. Front Immunol 9, 2831 (2018).

3. Tscharke, D. C., Croft, N. P., Doherty, P. C. & Gruta, N. L. L. Sizing up the key determinants of the CD8+ T cell response. Nat Rev Immunol 15, 705–716 (2015).

4. Kim, A. & Sadegh-Nasseri, S. Determinants of immunodominance for CD4 T cells. Curr Opin Immunol 34, 9–15 (2015).

5. Yewdell, J. W. Confronting Complexity: Real-World Immunodominance in Antiviral CD8+ T Cell Responses. Immunity 25, 533–543 (2006).

6. Mifsud, N. A. et al. Immunodominance Hierarchies and Gender Bias in Direct TCD8-*Cell* Alloreactivity. Am J Transplant 8, 121–132 (2008).

7. Schreiber, H., Wu, T. H., Nachman, J. & Kast, W. M. Immunodominance and tumor escape. Semin Cancer Biol 12, 25–31 (2002).

8. Crowe, S. R. et al. Differential Antigen Presentation Regulates the Changing Patterns of CD8+ T Cell Immunodominance in Primary and Secondary Influenza Virus Infections. J Exp Medicine 198, 399–410 (2003).

9. Kotturi, M. F. et al. Naive Precursor Frequencies and MHC Binding Rather Than the Degree of Epitope Diversity Shape CD8+ T Cell Immunodominance. J Immunol 181, 2124–2133 (2008).

10. Lazarski, C. A. et al. The Kinetic Stability of MHC Class II:Peptide Complexes Is a Key Parameter that Dictates Immunodominance. Immunity 23, 29–40 (2005).

11. Galea, I. et al. CD8+ T-cell cross-competition is governed by peptide–MHC class I stability. Eur J Immunol 42, 256–263 (2012).

12. Slifka, M. K. et al. Preferential Escape of Subdominant CD8+ T Cells During Negative Selection Results in an Altered Antiviral T Cell Hierarchy. J Immunol 170, 1231–1239 (2003).

13. Cukalac, T. et al. The Influenza Virus–Specific CTL Immunodominance Hierarchy in Mice Is Determined by the Relative Frequency of High-Avidity T Cells. J Immunol 192, 4061–4068 (2014).

14. Chen, W. & McCluskey, J. Immunodominance and Immunodomination: Critical Factors in Developing Effective CD8+ T-Cell–Based Cancer Vaccines. Adv Cancer Res 95, 203–247 (2006).

15. Kastenmuller, W. et al. Cross-competition of CD8+ T cells shapes the immunodominance hierarchy during boost vaccination. J Exp Medicine 204, 2187–2198 (2007).

16. Kedl, R. M., Schaefer, B. C., Kappler, J. W. & Marrack, P. T cells down-modulate peptide-MHC complexes on APCs in vivo. Nat Immunol 3, 27–32 (2002).

17. Lauron, E. J. et al. Cross-priming induces immunodomination in the presence of viral MHC class I inhibition. Plos Pathog 14, e1006883 (2018).

18. Waldman, A. D., Fritz, J. M. & Lenardo, M. J. A guide to cancer immunotherapy: from T cell basic science to clinical practice. Nat Rev Immunol 20, 651–668 (2020).

19. Gubin, M. M. et al. Checkpoint Blockade Cancer Immunotherapy Targets Tumour-Specific Mutant Antigens. Nature 515, 577–581 (2014).

20. Yadav, M. et al. Predicting immunogenic tumour mutations by combining mass spectrometry and exome sequencing. Nature 515, 572–576 (2014).

21. Castle, J. C. et al. Exploiting the Mutanome for Tumor Vaccination. Cancer Res. 72, 1081–1091 (2012).

22. Ott, P. A. et al. An immunogenic personal neoantigen vaccine for patients with melanoma. Nature 547, 217–221 (2017).

23. Sahin, U. et al. Personalized RNA mutanome vaccines mobilize poly-specific therapeutic immunity against cancer. Nature 547, 222–226 (2017).

24. Keskin, D. B. et al. Neoantigen vaccine generates intratumoral T cell responses in phase Ib glioblastoma trial. Nature 565, 234–239 (2019).

25. Rojas, L. A. et al. Personalized RNA neoantigen vaccines stimulate T cells in pancreatic cancer. Nature 1–7 (2023) doi:10.1038/s41586-023-06063-y.

26. Weber, J. S. et al. Individualised neoantigen therapy mRNA-4157 (V940) plus pembrolizumab versus pembrolizumab monotherapy in resected melanoma (KEYNOTE-942): a randomised, phase 2b study. Lancet (2024) doi:10.1016/s0140-6736(23)02268-7.

27. Palmer, C. D. et al. Individualized, heterologous chimpanzee adenovirus and self-amplifying mRNA neoantigen vaccine for advanced metastatic solid tumors: phase 1 trial interim results. Nat. Med. 28, 1619–1629 (2022).

28. Hu, Z., Ott, P. A. & Wu, C. J. Towards personalized, tumour-specific, therapeutic vaccines for cancer. Nat Rev Immunol 18, 168–182 (2018).

29. Paston, S. J., Brentville, V. A., Symonds, P. & Durrant, L. G. Cancer Vaccines, Adjuvants, and Delivery Systems. Front Immunol 12, 627932 (2021).

30. Hollingsworth, R. E. & Jansen, K. Turning the corner on therapeutic cancer vaccines. Npj Vaccines 4, 7 (2019).

31. Santos, P. M. & Butterfield, L. H. Dendritic Cell–Based Cancer Vaccines. J Immunol 200, 443–449 (2018).

32. Blass, E. & Ott, P. A. Advances in the development of personalized neoantigen-based therapeutic cancer vaccines. Nat Rev Clin Oncol 18, 215–229 (2021).

33. Beck, J. D. et al. mRNA therapeutics in cancer immunotherapy. Mol Cancer 20, 69 (2021).

34. D’Alise, A. Morena., et al. Phase I trial of viral vector based personalized vaccination elicits robust neoantigen specific antitumor T cell responses. Clin. Cancer Res. 30, 2412–2423 (2024).

35. Lopez, J. et al. Autogene cevumeran with or without atezolizumab in advanced solid tumors: a phase 1 trial. Nat. Med. 1–13 (2025) doi:10.1038/s41591-024-03334-7.

36. Kranz, L. M. et al. Systemic RNA delivery to dendritic cells exploits antiviral defence for cancer immunotherapy. Nature 534, 396–401 (2016).

37. Kreiter, S. et al. Mutant MHC class II epitopes drive therapeutic immune responses to cancer. Nature 520, 692–696 (2015).

38. Sethna, Z. et al. RNA neoantigen vaccines prime long-lived CD8+ T cells in pancreatic cancer. Nature 1–10 (2025) doi:10.1038/s41586-024-08508-4.

39. Yadav, M. et al. Predicting immunogenic tumour mutations by combining mass spectrometry and exome sequencing. Nature 515, 572–576 (2014).

40. Capietto, A.-H. et al. Mutation position is an important determinant for predicting cancer neoantigens. J Exp Med 217, e20190179 (2020).

41. Vesin, B. et al. Optimized lentiviral backbone induces robust and diverse T cell immunity against neoantigens to counteract tumor heterogeneity. npj Vaccines 10, 143 (2025).

42. Kranz, L. M. et al. Systemic RNA delivery to dendritic cells exploits antiviral defence for cancer immunotherapy. Nature 534, 396–401 (2016).

43. Peters, D. et al. RNA-encoded Interleukin 2 with Extended Bioavailability Amplifies RNA Vaccine–Induced Antitumor T-cell Immunity. Cancer Immunol. Res. 12, 1409–1420 (2024).

44. Rees, W. et al. An inverse relationship between T cell receptor affinity and antigen dose during CD4 + T cell responses in vivo and in vitro. Proc National Acad Sci 96, 9781–9786 (1999).

45. Jamieson, B. D. & Ahmed, R. T cell memory. Long-term persistence of virus-specific cytotoxic T cells. J Exp Medicine 169, 1993–2005 (1989).

46. Sant, A. J. et al. The relationship between immunodominance, DM editing, and the kinetic stability of MHC class II:peptide complexes. Immunol Rev 207, 261–278 (2005).

47. Tung, J. & Sant, A. J. Orchestration of CD4 T Cell Epitope Preferences after Multipeptide Immunization. J Immunol 191, 764–772 (2013).

48. Weaver, J. M., Chaves, F. A. & Sant, A. J. Abortive activation of CD4 T cell responses during competitive priming in vivo. Proc National Acad Sci 106, 8647–8652 (2009).

49. Kedl, R. M. et al. T Cells Compete for Access to Antigen-Bearing Antigen-Presenting Cells. J Exp Medicine 192, 1105–1114 (2000).

50. Gober, J. G. et al. MHC2-SCALE enhances identification of immunogenic neoantigens. iScience 28, 112212 (2025).

51. Harndahl, M. et al. Peptide-MHC class I stability is a better predictor than peptide affinity of CTL immunogenicity. Eur J Immunol 42, 1405–1416 (2012).

52. Strønen, E. et al. Targeting of cancer neoantigens with donor-derived T cell receptor repertoires. Science 352, 1337–1341 (2016).

53. Burger, M. L. et al. Antigen dominance hierarchies shape TCF1+ progenitor CD8 T cell phenotypes in tumors. Cell 184, 4996–5014.e26 (2021).

54. Hellman, L. M. et al. Differential scanning fluorimetry based assessments of the thermal and kinetic stability of peptide–MHC complexes. J Immunol Methods 432, 95–101 (2016).

55. Anderson, B. R. et al. Incorporation of pseudouridine into mRNA enhances translation by diminishing PKR activation. Nucleic Acids Res 38, 5884–5892 (2010).

56. Anderson, B. R. et al. Nucleoside modifications in RNA limit activation of 2′-5′-oligoadenylate synthetase and increase resistance to cleavage by RNase L. Nucleic Acids Res 39, 9329–9338 (2011).

57. Krienke, C. et al. A noninflammatory mRNA vaccine for treatment of experimental autoimmune encephalomyelitis. Science 371, 145–153 (2021).

58. Schnell, A. et al. Targeting PGLYRP1 promotes antitumor immunity while inhibiting autoimmune neuroinflammation. Nat. Immunol. 24, 1908–1920 (2023).

59. Castiglioni, A. et al. Combined PD-L1/TGFβ blockade allows expansion and differentiation of stem cell-like CD8 T cells in immune excluded tumors. Nat. Commun. 14, 4703 (2023).

60. Hudson, W. H. et al. Proliferating Transitory T Cells with an Effector-like Transcriptional Signature Emerge from PD-1+ Stem-like CD8+ T Cells during Chronic Infection. Immunity 51, 1043–1058.e4 (2019).

61. Blaser, C. et al. β-Galactoside-binding protein secreted by activated T cells inhibits antigen-induced proliferation of T cells. Eur. J. Immunol. 28, 2311–2319 (1998).

62. Bakos, E. et al. CCR2 Regulates the Immune Response by Modulating the Interconversion and Function of Effector and Regulatory T Cells. J. Immunol. 198, 4659–4671 (2017).

63. Marçais, A. et al. Cell-Autonomous CCL5 Transcription by Memory CD8 T Cells Is Regulated by IL-4. J. Immunol. 177, 4451–4457 (2006).

64. D’Alise, A. M. et al. Adenoviral-based vaccine promotes neoantigen-specific CD8+ T cell stemness and tumor rejection. Sci Transl Med 14, eabo7604 (2022).

65. Zhao, X., Shan, Q. & Xue, H.-H. TCF1 in T cell immunity: a broadened frontier. Nat. Rev. Immunol. 22, 147–157 (2022).

66. Escobar, G., Mangani, D. & Anderson, A. C. T cell factor 1: A master regulator of the T cell response in disease. Sci. Immunol. 5, (2020).

67. Levin, N. et al. Neoantigen-specific stimulation of tumor-infiltrating lymphocytes enables effective TCR isolation and expansion while preserving stem-like memory phenotypes. J. Immunother. Cancer 12, e008645 (2024).

68. Tahtinen, S. et al. IL-1 and IL-1ra are key regulators of the inflammatory response to RNA vaccines. Nat Immunol 1–11 (2022) doi:10.1038/s41590-022-01160-y.

69. Beltra, J.-C. et al. Developmental Relationships of Four Exhausted CD8+ T Cell Subsets Reveals Underlying Transcriptional and Epigenetic Landscape Control Mechanisms. Immunity 52, 825–841.e8 (2020).

70. Kaseke, C. et al. HLA class-I-peptide stability mediates CD8+ T cell immunodominance hierarchies and facilitates HLA-associated immune control of HIV. Cell Reports 36, 109378 (2021).

71. Lineburg, K. E. et al. CD8+ T cells specific for an immunodominant SARS-CoV-2 nucleocapsid epitope cross-react with selective seasonal coronaviruses. Immunity 54, 1055–1065.e5 (2021).

72. Galea, I. et al. CD8+ T-cell cross-competition is governed by peptide–MHC class I stability. Eur J Immunol 42, 256–263 (2012).

73. Chen, L. et al. B7-H1 maintains the polyclonal T cell response by protecting dendritic cells from cytotoxic T lymphocyte destruction. Proc National Acad Sci 115, 201722043 (2018).

74. Rytelewski, M. et al. Suppression of Immunodominant Antitumor and Antiviral CD8+ T Cell Responses by Indoleamine 2,3-Dioxygenase. Plos One 9, e90439 (2014).

75. Rodriguez, F., Harkins, S., Slifka, M. K. & Whitton, J. L. Immunodominance in Virus-Induced CD8 + T-Cell Responses Is Dramatically Modified by DNA Immunization and Is Regulated by Gamma Interferon. J Virol 76, 4251–4259 (2002).

76. Haeryfar, S. M. M., DiPaolo, R. J., Tscharke, D. C., Bennink, J. R. & Yewdell, J. W. Regulatory T Cells Suppress CD8+ T Cell Responses Induced by Direct Priming and Cross-Priming and Moderate Immunodominance Disparities. J Immunol 174, 3344–3351 (2005).

77. Thirdborough, S. M. et al. Tapasin shapes immunodominance hierarchies according to the kinetic stability of peptide – MHC class I complexes. Eur J Immunol 38, 364–369 (2008).

78. Kedl, R. M. et al. T Cells Compete for Access to Antigen-Bearing Antigen-Presenting Cells. J Exp Medicine 192, 1105–1114 (2000).

79. Sandberg, J. K., et al. Superdominance among immunodominant H-2Kb-restricted epitopes and reversal by dendritic cell-mediated antigen delivery. J. Immunol. (Baltim., Md : 1950) 160, 3163–9 (1998).

80. Kedl, R. M., Schaefer, B. C., Kappler, J. W. & Marrack, P. T cells down-modulate peptide-MHC complexes on APCs in vivo. Nat Immunol 3, 27–32 (2002).

81. Rodriguez, F., Harkins, S., Slifka, M. K. & Whitton, J. L. Immunodominance in Virus-Induced CD8 + T-Cell Responses Is Dramatically Modified by DNA Immunization and Is Regulated by Gamma Interferon. J Virol 76, 4251–4259 (2002).

82. Zhang, S.-Q. et al. High-throughput determination of the antigen specificities of T cell receptors in single cells. Nat Biotechnol 36, 1156–1159 (2018).

83. Marzo, A. L. et al. Initial T cell frequency dictates memory CD8+ T cell lineage commitment. Nat Immunol 6, 793–799 (2005).

84. Liu, F., Whitton, J. L. & Slifka, M. K. The Rapidity with Which Virus-Specific CD8+ T Cells Initiate IFN-γ Synthesis Increases Markedly over the Course of Infection and Correlates with Immunodominance. J Immunol 173, 456–462 (2004).

85. Galassi, C., Chan, T. A., Vitale, I. & Galluzzi, L. The hallmarks of cancer immune evasion. Cancer Cell 42, 1825–1863 (2024).

86. Flecken, T. et al. Immunodominance and functional alterations of tumor-associated antigen-specific CD8+ T-cell responses in hepatocellular carcinoma. Hepatology 59, 1415–1426 (2014).

87. Keane, C. et al. The T-cell Receptor Repertoire Influences the Tumor Microenvironment and Is Associated with Survival in Aggressive B-cell Lymphoma. Clin. Cancer Res. 23, 1820–1828 (2017).

88. Postow, M. A. et al. Peripheral T cell receptor diversity is associated with clinical outcomes following ipilimumab treatment in metastatic melanoma. J. Immunother. Cancer 3, 23 (2015).

89. Kansy, B. A. et al. T cell receptor richness in peripheral blood increases after cetuximab therapy and correlates with therapeutic response. OncoImmunology 7, e1494112 (2018).

90. Jia, Q. et al. Diversity index of mucosal resident T lymphocyte repertoire predicts clinical prognosis in gastric cancer. OncoImmunology 4, e1001230 (2015).

91. Pothuri, V. S. et al. Intratumoral T-cell receptor repertoire composition predicts overall survival in patients with pancreatic ductal adenocarcinoma. OncoImmunology 13, 2320411 (2024).

92. Rodenko, B. et al. Generation of peptide–MHC class I complexes through UV-mediated ligand exchange. Nat Protoc 1, 1120–1132 (2006).

93. Toebes, M. et al. Design and use of conditional MHC class I ligands. Nat Med 12, 246–251 (2006).

94. Capietto, A.-H. et al. Mutation position is an important determinant for predicting cancer neoantigens. J Exp Med 217, e20190179 (2020).

95. Jenkins, M. K. & Moon, J. J. The Role of Naive T Cell Precursor Frequency and Recruitment in Dictating Immune Response Magnitude. J Immunol 188, 4135–4140 (2012).

96. Obar, J. J., Khanna, K. M. & Lefrançois, L. Endogenous Naive CD8+ T Cell Precursor Frequency Regulates Primary and Memory Responses to Infection. Immunity 28, 859–869 (2008).

97. Legoux, F. P. & Moon, J. J. Peptide:MHC Tetramer-based Enrichment of Epitope-specific T cells. J. Vis. Exp. (2012) doi:10.3791/4420.

98. Gaublomme, J. T. et al. Nuclei multiplexing with barcoded antibodies for single-nucleus genomics. Nat. Commun. 10, 2907 (2019).

99. Hao, Y. et al. Dictionary learning for integrative, multimodal and scalable single-cell analysis. Nat. Biotechnol. 42, 293–304 (2024).

